# Detecting and quantifying heterogeneity in susceptibility using contact tracing data

**DOI:** 10.1101/2023.10.04.560944

**Authors:** Beth M. Tuschhoff, David A. Kennedy

## Abstract

The presence of heterogeneity in susceptibility, differences between hosts in their likelihood of becoming infected, can fundamentally alter disease dynamics and public health responses, for example, by changing the final epidemic size, the duration of an epidemic, and even the vaccination threshold required to achieve herd immunity. Yet, heterogeneity in susceptibility is notoriously difficult to detect and measure, especially early in an epidemic. Here we develop a method that can be used to detect and estimate heterogeneity in susceptibility given contact by using contact tracing data, which is typically collected early in the course of an outbreak. This approach provides the capability, given sufficient data, to estimate and account for the effects of this heterogeneity before they become apparent during an epidemic. It additionally provides the capability to analyze the wealth of contact tracing data available for previous epidemics and estimate heterogeneity in susceptibility for disease systems in which it has never been estimated previously. The premise of our approach is that highly susceptible individuals become infected more often than less susceptible individuals, and so individuals not infected after appearing in contact networks should be less susceptible than average. This change in susceptibility can be detected and quantified when individuals show up in a second contact network after not being infected in the first. To develop our method, we simulated contact tracing data from artificial populations with known levels of heterogeneity in susceptibility according to underlying discrete or continuous distributions of susceptibilities. We analyzed this data to determine the parameter space under which we are able to detect heterogeneity and the accuracy with which we are able to estimate it. We found that our power to detect heterogeneity increases with larger sample sizes, greater heterogeneity, and intermediate fractions of contacts becoming infected in the discrete case or greater fractions of contacts becoming infected in the continuous case. We also found that we are able to reliably estimate heterogeneity and disease dynamics. Ultimately, this means that contact tracing data alone is sufficient to detect and quantify heterogeneity in susceptibility.

## 1. Introduction

At the outset of an epidemic, public health responses depend on estimates of the final epidemic size, the peak number of cases, the timing of the peak, and the herd immunity threshold. Compartmental models such as the susceptible-infected-recovered (SIR) model are commonly used to model infectious disease dynamics and predict outcomes, but there are limitations to this approach (Keeling and Danon, 2009; Roberts et al., 2015; Tolles and Luong, 2020; Dhar, 2020). Namely, SIR models tend to oversimplify the complexity of disease dynamics, resulting in discrepancies between the model predictions and epidemic data (Keeling and Danon, 2009). One of the simplifying assumptions of the standard SIR model is that all host individuals are the same. However, this is often false: individuals can be heterogeneous in many ways (Woolhouse et al., 1997; VanderWaal and Ezenwa, 2016) including with regard to their likelihood of becoming infected, hereafter referred to as heterogeneity in susceptibility (Dwyer et al., 1997).

Heterogeneity in susceptibility can have a large impact on infectious disease dynamics (Dwyer et al., 1997; Gomes et al., 2014; Langwig et al., 2017; Gomes et al., 2022). Increased amounts of heterogeneity in susceptibility result in a lower peak number of cases, different timing of the peak, smaller final epidemic size, and lower herd immunity thresholds (Aguas et al., 2020; Gomes et al., 2022; Montalbán et al., 2022). As a result, disease control programs (Anderson and May, 1984) and epidemiological models (Dwyer et al., 1997; Langwig et al., 2017; King et al., 2018; Gomes et al., 2019) may need to account for heterogeneity in susceptibility if they are to be optimally useful. Accurate early predictions of disease dynamics could give policy makers critical information to make decisions, but heterogeneity in susceptibility is notoriously difficult to measure (Elderd et al., 2008). Moreover, the effects of heterogeneity in susceptibility are typically small during the earliest phases of epidemics and only become apparent later, making it even more challenging to estimate heterogeneity in susceptibility in real time and account for its effects. It would therefore be useful to develop new methods for quantifying the degree of heterogeneity in host susceptibility early in epidemics.

Existing methods to quantify heterogeneity in susceptibility are not adequate for estimation in real time because they rely on using data that is either collected later in epidemics or that typically cannot be collected due to ethical or logistical constraints. Dwyer et al. (1997), Ben-Ami et al. (2010), and Langwig et al. (2017) used laboratory dose-response and field transmission experiments to estimate heterogeneity in susceptibility, but these experimental methods are not feasible for application in real time or for human epidemics in general due to time constraints and ethical concerns. Gomes et al. (2019) compared disease incidence across municipalities in several countries to construct Lorenz curves and fit susceptibility risk distributions, but this method requires a substantial amount of data that would not be available early in an epidemic. Smith et al. (2005) and Corder et al. (2020) used morbidity data to fit models and estimate heterogeneity, but this method cannot be used until later in an epidemic when there is sufficient data to fit curves. Gomes et al. (2022) also used curve fitting with mortality data that could be implemented once at least four months of data were available, but their method is heavily dependent on the underlying model and assumptions. With the recent increased interest in real-time estimation, Anderson et al. (2023) developed a method to estimate within-household heterogeneity in susceptibility, but this is not the same as the population-level heterogeneity that drives population-level disease dynamics. Here we develop a novel method to identify and quantify host heterogeneity in susceptibility using contact tracing data, which can be collected early in an epidemic. Contact tracing is often performed to mitigate the spread of pathogens that are otherwise difficult to control (Eames and Keeling, 2003; Hossain et al., 2022), and therefore, our method should not require the collection of any data beyond that which would already be collected for other purposes.

Contact tracing typically takes one of two forms: forward and backward. Forward contact tracing attempts to find all the contacts of an infected person to whom the disease could transmit. This is done by identifying infected individuals and all their known contacts. The contacts are then quarantined and monitored for disease. For any contact that is infected, the process is repeated with their contacts. Backward contact tracing attempts to identify the contact of an infected person from whom the disease transmitted. In practice, both methods can be employed simultaneously in an effort to maximize the effectiveness of contact tracing efforts (Bradshaw et al., 2021), and the data on infected individuals and their contacts are typically recorded. When done thoroughly, contact tracing data provide information about the infection status of individuals that have been in contact with an infected individual. As we will explain, when contact tracing data tracks specific individuals through multiple exposure events, it can be used to quantify heterogeneity in susceptibility given contact through the method that we develop here.

Our method uses the fact that average susceptibility decreases over time in a population with heterogeneity in susceptibility (Fig 1). This is because individuals with high susceptibility are more likely to be infected than individuals with low susceptibility for a given exposure level. Individuals that show up in a second contact tracing network, after not being infected in the first, should therefore have a lower risk of infection than individuals that show up in a network for the first time. In the rest of the paper, we establish our method and analyze its effectiveness for two cases: a population with two discrete susceptibility levels and a population with continuous variation in susceptibility. Notably, the selection of these two cases is arbitrary, and our method is flexible enough that it could be employed for any distribution of heterogeneity in susceptibility.

**Figure 1:**
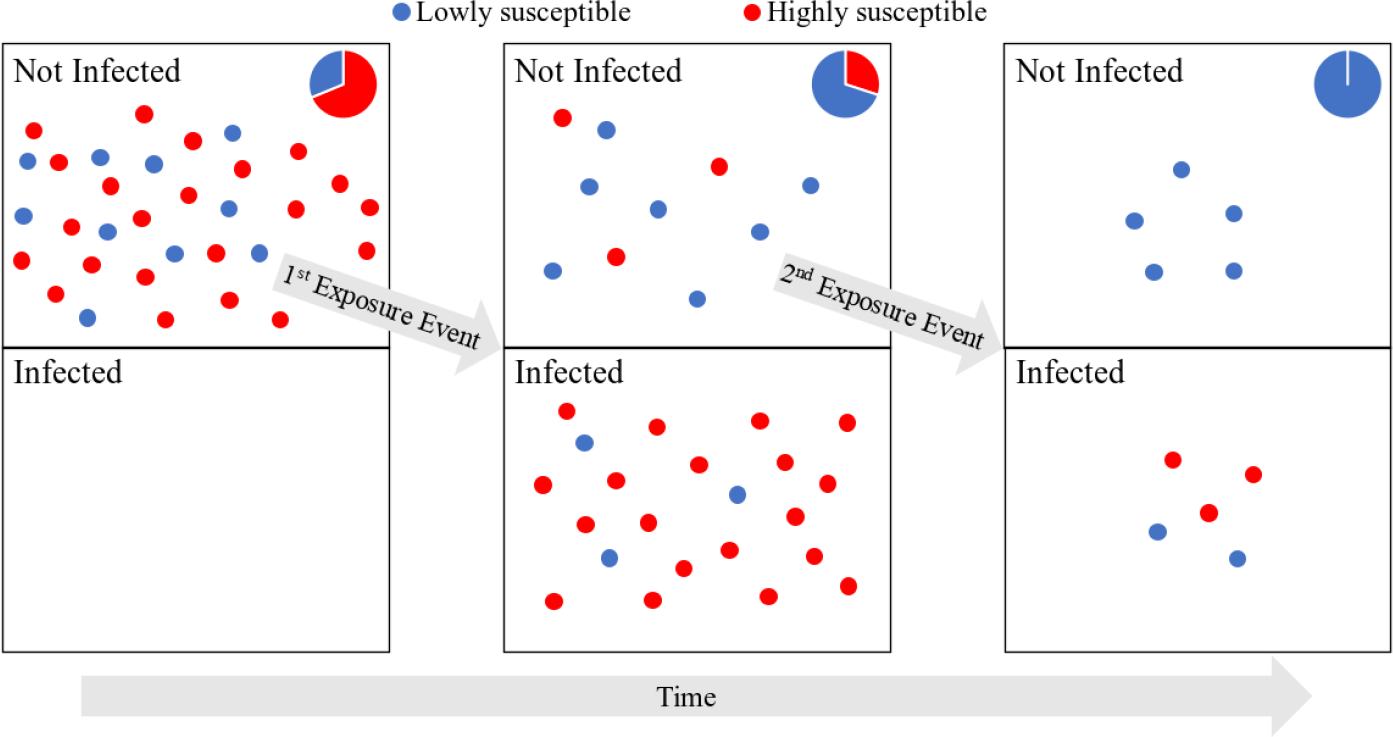
Average susceptibility decreases over exposure events in a heterogeneous population. The figure depicts individuals infected and not infected over two exposure events in a heterogeneous population with more susceptible (red) and less susceptible (blue) individuals. The pie charts show the composition of the not infected population. Average susceptibility in the not infected population decreases after each exposure event as the more susceptible individuals are primarily infected. Note that if the population lacked heterogeneity in susceptibility, all individuals would be either red or blue, and thus, susceptibility would not change.

## 2. Methods and Results

Our method to detect and quantify heterogeneity in susceptibility exploits the change in average susceptibility over multiple exposure events that would be expected to occur if a population had heterogeneity in susceptibility (Fig 1). Given contact with an infectious individual, individuals with high susceptibility are more likely to be infected than those with low susceptibility. This creates a selection process in which susceptibility should on average decline in a heterogeneous host population following each exposure event. This change in average susceptibility provides a way to identify and estimate the level of heterogeneity early in an epidemic despite the seemingly small effects of heterogeneity at the beginning of epidemics. Notably, no change in average susceptibility should occur in a population that lacks heterogeneity in susceptibility.

This method employs contact tracing data. With contact tracing data, there are multiple contact networks that are each composed of an infected individual and the known contacts the infected individual had during their infectious period. This means each contact network is a set of exposure events where contacts are exposed to a pathogen and have a chance of being infected. In order for our method to work, there must be individuals that show up in at least two separate contact networks such that they are exposed but not infected in the first of these networks. At the start of the second exposure event, these individuals would have been previously exposed but not infected (henceforth called focal individuals). This contact network must also contain naive contacts: individuals that have not been previously exposed to the pathogen. The basis of our method is to compare the fraction of naive individuals and focal individuals that become infected in the second contact network; if there is no heterogeneity in susceptibility, focal individuals should have the same susceptibility as naive individuals, whereas if there is heterogeneity in susceptibility, focal individuals should on average be less susceptible than naive individuals. This difference in susceptibility arises due to the selection process for infection of more susceptible individuals (Fig 1).

To compute the number of naive and focal individuals infected, there must be data on which specific individuals are infected and which individuals are showing up in a contact network for a second time, which would be available for example if individuals were identifiable between contact networks. There must also be a sufficient sample size to detect heterogeneity in susceptibility. Here we explore the effects of sample size, level of heterogeneity, and infection probability on our ability to detect and quantify heterogeneity in susceptibility.

We apply this method to two underlying models describing the distribution of individuals’ susceptibilities. In one underlying model (discrete case), it is assumed that the population is composed of two host types where each host type has a different susceptibility or probability of being infected given contact. Discrete susceptibility types might be expected when heterogeneity in susceptibility is predominantly accounted for by a small number of factors that create groups in the population with distinct susceptibilities. For example, genetic polymorphisms could be selected for that increase resistance to a pathogen, resulting in populations containing a mixture of individuals with and without the mutation such as was seen for HIV (Huang et al., 1996). Likewise, prior exposure, whether natural or vaccine-induced, to a pathogen or related pathogen could create more resistant subpopulations such as with milkmaids not developing smallpox after contracting cowpox (Barquet and Domingo, 1997). Behaviors like handwashing and mask wearing (Larson, 1999; Van der Sande et al., 2008) or host nutritional status (Chandra, 1979) could also produce approximately binary outcomes for susceptibility to infection.

In the other underlying model (continuous case), it is assumed that the population is composed of hosts with a continuous range of susceptibilities such that each host’s probability of being infected given contact is unique. This situation might be expected when there is a complex combination of factors dictating heterogeneity in susceptibility or when the cause of heterogeneity is a trait that continuously varies across individuals. For instance, variability in gene expression, which could be affected by epigenetics, copy number variations, and sequence polymorphisms, is associated with disease susceptibility (Li et al., 2010). In addition, some of the factors that lead to discrete variation in susceptibility could also have a continuous effect such as the degree of cleanliness achieved by handwashing (Larson, 1999) or continuous variation in nutrients. Beyond a complex combination of factors, there could also be situations where a continuously varying trait like body mass (Dobner and Kaser, 2018) or the level of antibodies induced in an immune response (Plotkin, 2008) explains the heterogeneity in susceptibility in the population.

### 2.1. Methods

Our method is comprised of two parts: detecting heterogeneity in susceptibility and quantifying it if present. The former is a hypothesis testing problem, and the latter is a parameter estimation problem. For the detection of heterogeneity, we test the hypothesis that there is heterogeneity in susceptibility against the null hypothesis that there is homogeneity in susceptibility.

#### 2.1.1. Detection of heterogeneity in susceptibility

We consider *F* contact networks that each contain *N*_*i*_ − 1 naive individuals and one focal individual where *i* is the set of contact networks. For simplicity, we assume *N*_*i*_ are equal for all *i* and thus drop the subscript. Note that this assumption can be easily relaxed. We therefore have a total of *F* (*N*− 1) naive individuals and *F* focal individuals. We first compute the fractions of naive, focal, and total individuals infected. The fractions of naive and focal individuals infected are estimates for the probability of a naive or focal individual being infected (*p*_*n*_ and *p*_*f*_ respectively). The fraction of total individuals infected is an estimate for the average probability of being infected 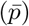. We then calculate the log-likelihood of the data (numbers of individuals infected) under each hypothesis as a sum of the log-likelihoods for the number of each type of individual infected where

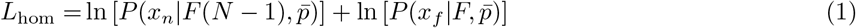

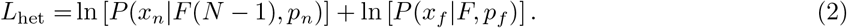

*L*_hom_ is the log-likelihood of the data under the null hypothesis that there is homogeneity in susceptibility, so we assume all individuals have the same probability of being infected, regardless of their number of exposures to the pathogen 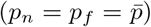. *L*_het_ is the log-likelihood under the alternative hypothesis that there is heterogeneity in susceptibility, so we assume naive and focal individuals have different probabilities of being infected due to the infection selection process that occurs when heterogeneity is present (*p*_*n*_ ≠ *p*_*f*_). These log-likelihoods are calculated identically regardless of whether the heterogeneity is discrete or continuous. *P* (*x*| *n, p*) is the probability of observing *x* individuals infected out of *n* individuals exposed with probability *p* of being infected and is distributed according to a binomial distribution. The number of naive individuals infected has distribution Binom(*n* = *F* (*N* − 1), *p*_*n*_), and the number of focal individuals infected has distribution Binom(*n* = *F, p*_*f*_). *x*_*n*_ and *x*_*f*_ are the numbers of naive and focal individuals infected respectively where 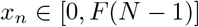 and *x*_*f*_ *∈* [0, *F*]. *p*_*n*_, *p*_*f*_ and 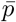are estimated from the data as 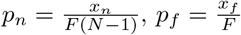, and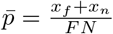. The log-likelihoods of the data under each hypothesis were compared using a likelihood ratio test with one degree of freedom and significance level *α* = 0.05.

Here, we simulated data to test our method. To do so, we first set parameters dictating the sample size and heterogeneity present in the population. Then, we simulated initial exposure events with *N* individuals in each network and kept uninfected individuals as our focal individuals. For each focal individual, we then simulated a second exposure event with that focal individual and *N*− 1 naive individuals. The susceptibilities of the naive individuals were drawn randomly from the same heterogeneity distribution set for the starting population. We recorded the fraction of each type of individual (i.e. focal or naive) infected in the second exposure events and calculated the log-likelihood of the simulated data under our two hypotheses. Then, we compared the hypotheses with a likelihood ratio test. We ran 1, 000 simulations for each set of parameters to determine our statistical power to detect heterogeneity in susceptibility with that parameter combination. All simulations and data analysis were performed in R version 4.0.3 (R Core Team, 2020).

For the discrete case, we simulated data using two types of individuals (denoted *A* and *B*), but we note that the aforementioned factors could potentially be combined to result in more than two distinct groupings, and similar methods could be applied for these situations. At the beginning of each simulation, we set the probability of being infected for each type of individual, *p*_*A*_ and *p*_*B*_, where *p*_*A*_ *∈* [0, 1] and *p*_*B*_ *∈* [0, *p*_*A*_]. We also set the fraction of the starting population that is type *A* (*f*_*A*_) where *f*_*A*_ *∈* [0, 1]. All three parameters *p*_*A*_, *p*_*B*_, and *f*_*A*_ affect the level of heterogeneity in susceptibility in the population.

We later calculated the coefficient of variation of the risk of being infected for this discrete case (*C*_*d*_) and the expected fraction of naive individuals infected (*E*_*d*_) from *p*_*A*_, *p*_*B*_, and *f*_*A*_ to better summarize the results. The risks of being infected for type *A* and *B* individuals, *r*_*A*_ and *r*_*B*_ respectively, are shown below. These equations are derived from the formula for the probability of being infected 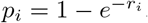, *i* = *A, B*.

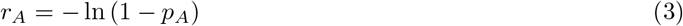

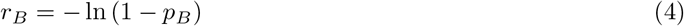

The coefficient of variation is defined as the standard deviation divided by the mean. Hence, *C*_*d*_ is the standard deviation of risk divided by the mean risk (Supplementary information S1) and is given by

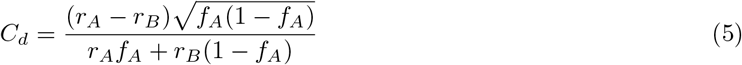

*E*_*d*_ is the same as the mean probability of being infected 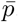, which is given by

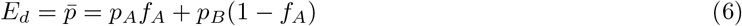

We additionally defined the sample size for the simulation by setting the number of individuals in each exposure group *N* and the number of focal individuals *F*. For our simulations, we used *N* = 5 and *F* = 50 or 200.

For the continuous case, in contrast to the discrete case just discussed, each individual in the population has a different risk of being infected. Here, we assume that individuals’ risks for being infected follow a gamma distribution, but as in the discrete case, other distributions could be used. We chose to use a gamma distribution for illustration purposes because it is flexible and has been used to model heterogeneous populations previously (Dwyer et al., 1997; Langwig et al., 2017).

At the beginning of each simulation, we set the parameters *k* and *θ*, respectively the shape and scale of the gamma distribution, that dictate the risk distribution where *k, θ >* 0. For ease of interpretation, we present our results with respect to the coefficient of variation of risk for continuous variation *C*_*c*_ and expected fraction of naive individuals infected *E*_*c*_. As in the discrete case, the risk *r*_*i*_ for the *i*th individual being infected is related to the probability of being infected such that 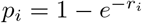and thus

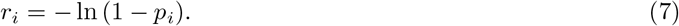

As it is gamma distributed, the risk distribution has standard deviation 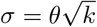can be simplified to and mean *μ* = *kθ*. So, *C*_*c*_ can be simplified to

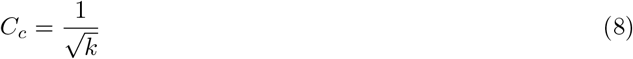

*E*_*c*_ is the same as the mean probability of being infected 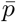 and is derived in Dwyer et al. (1997) as

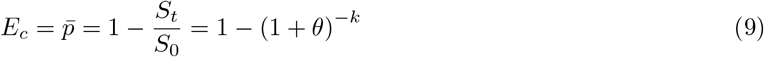

where *S*_0_ and *S*_*t*_ are the number of susceptible individuals at the beginning and end of an exposure round respectively.

We additionally defined the sample size for the simulation by setting the number of individuals in each exposure group *N* and the number of focal individuals *F*. As in the discrete case, we use *N* = 5 and *F* = 50 or 200.

We tested the ability of our method to detect heterogeneity in susceptibility for each potential combination of *f*_*A*_, *F, C*_*d*_ *∈* [0, 3] with step size 0.02, and *E*_*d*_ *∈* [0.02, 0.98] with step size 0.02 in the discrete case and *F, C*_*c*_ *∈* [0, 3] with step size 0.02, and *E*_*c*_ *∈* [0.02, 0.98] with step size 0.02 in the continuous case. This was done for 1, 000 simulations to compute the statistical power of the method. We did not simulate *E*_*d*_ = 0, 1 or *E*_*c*_ = 0, 1 because such values preclude heterogeneity in susceptibility. We examined *C*_*d*_, *C*_*c*_ *∈* [0, 3] because this captures most of the range of published values for the coefficient of variation of risk we could find: 0.0007 to 3.33 (Dwyer et al., 1997, 2000; Smith et al., 2005; Ben-Ami et al., 2008; Elderd et al., 2008; Ben-Ami et al., 2010; Pessoa et al., 2014; Langwig et al., 2017; King et al., 2018; Gomes et al., 2019; Corder et al., 2020; Gomes et al., 2022).

#### 2.1.2. Quantification of heterogeneity in susceptibility

Given the detection of heterogeneity in susceptibility, the next question is whether that heterogeneity will substantially impact disease dynamics. To determine whether it will, we need to ask whether contact tracing data is sufficient to estimate the parameters of SIR models that include heterogeneity in susceptibility and whether those parameter estimates accurately capture disease dynamics. To do so, we fit the parameters of our underlying risk distributions using simulated contact tracing data as above. Parameter values used to simulate the contact tracing data for the discrete and continuous heterogeneity cases are provided in Table 1.

**Table 1:**
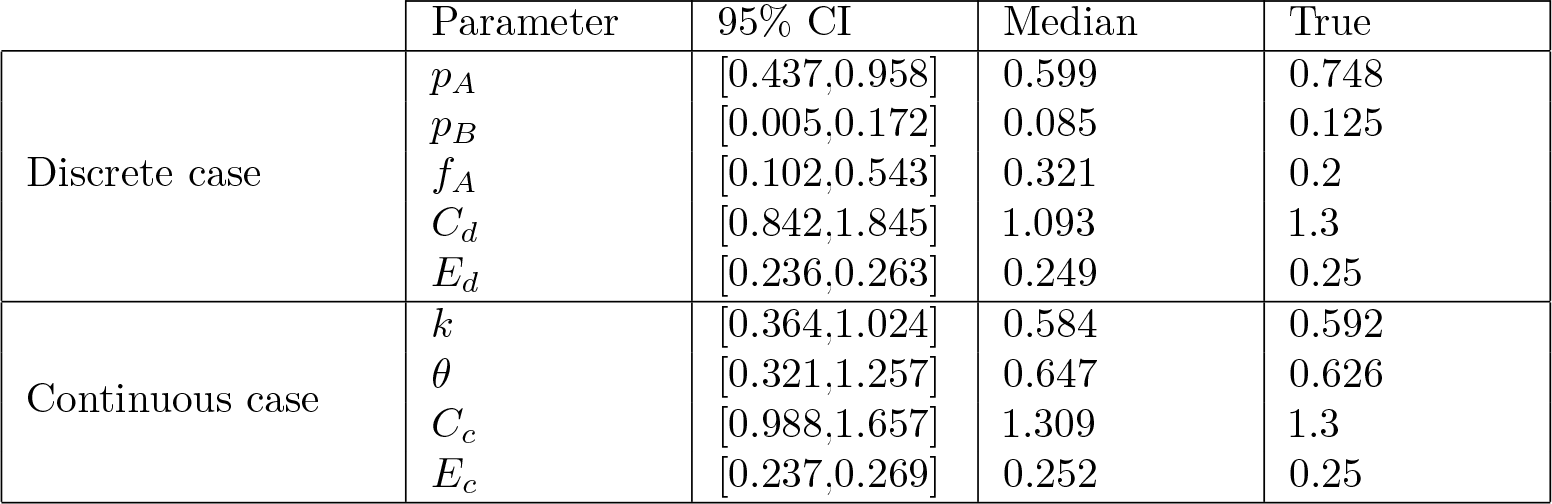
The 95% CIs, medians, and true values for parameters estimated from MCMC in the discrete and continuous cases with *F* = 1000 and *N* = 5.

|  | Parameter | 95% CI | Median | True |
| --- | --- | --- | --- | --- |
| Discrete case | $p_A$ | [0.437,0.958] | 0.599 | 0.748 |
| | $p_B$ | [0.005,0.172] | 0.085 | 0.125 |
| | $f_A$ | [0.102,0.543] | 0.321 | 0.2 |
| | $C_d$ | [0.842,1.845] | 1.093 | 1.3 |
| | $E_d$ | [0.236,0.263] | 0.249 | 0.25 |
| Continuous case | $k$ | [0.364,1.024] | 0.584 | 0.592 |
| | $\theta$ | [0.321,1.257] | 0.647 | 0.626 |
| | $C_c$ | [0.988,1.657] | 1.309 | 1.3 |
| | $E_c$ | [0.237,0.269] | 0.252 | 0.25 |

We generated posterior distributions for both models using Metropolis-Hastings MCMC. In the discrete case, our MCMC chain had length 30,000,000 with a burn-in of 15,000,000 and thinning interval 1,500. For all three parameters, we used flat priors and uniform proposal distributions. Our proposal distributions were *p*_*A*_*∼* Unif(0, 1), *p*_*B*_ *∼*Unif(0, *p*_*A*_), and *f*_*A*_ *∼*Unif(0, 1). There is not a simple, analytic likelihood function for the likelihood of the data given a proposed parameter set, so the likelihood was estimated by simulation with Approximate Bayesian Computation (ABC), where the likelihood estimate was determined by comparing the fraction of simulations that provided results that were within a pre-specified error tolerance of the actual data (Beaumont et al., 2002). To do so, we ran 100 simulations of the number of focal and naive individuals infected across *F* contact networks for a proposed parameter set. We then calculated the fraction of simulations where the number of individuals infected was within a 1% error tolerance of the number infected in the true data. Note that our results are fairly insensitive to this error tolerance (Supplementary information S4). This simulation was done separately for focal and naive individuals. We then computed the overall log-likelihood as a sum of the logs of those fractions. We assessed convergence of the chains by visually inspecting the resulting trace plots and marginal posterior distributions for each parameter. In the continuous case, our MCMC chain had length 600,000 with a burn-in of 200,000 and thinning interval 100. We used an exponential prior Exp(2) for *k* because known values of *C*_*c*_ suggest that *k* is likely to be small (Dwyer et al., 1997, 2000; Smith et al., 2005; Ben-Ami et al., 2008; Elderd et al., 2008; Ben-Ami et al., 2010; Pessoa et al., 2014; Langwig et al., 2017; King et al., 2018; Gomes et al., 2019; Corder et al., 2020; Gomes et al., 2022). We used a flat prior for *θ* for all values [0, inf) and a multivariate lognormal proposal distribution (*k, θ*) *∼* MLogNorm 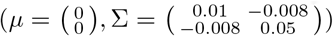. We assessed convergence of the chains by visually inspecting the resulting trace plots and marginal posterior distributions for each parameter (Kennedy et al., 2015).

We then used these parameter estimates to generate SIR dynamics. Notably, the system of differential equations describing the discrete and continuous cases differ. For the discrete case, we implemented the following system of ordinary differential equations:

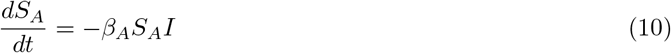

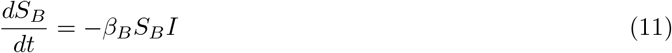

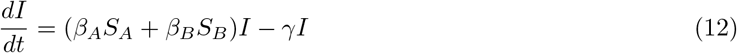

*S*_*A*_ and *S*_*B*_ are the susceptible individuals of types *A* and *B*, and *I* is the infected individuals where *I* includes infected *A* and infected *B* individuals such that *I* = *I*_*A*_ + *I*_*B*_. At the start of each SIR simulation, we determine the fraction of the population to allocate as *A* and *B* from *f*_*A*_. We also set the basic reproduction number 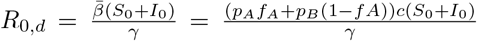 at an assumed “true” value where 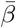 is the average transmission rate and *S*_0_ + *I*_0_ is the population size. *R*_0,*d*_ is often a reasonably well approximated value, and it does not change with heterogeneity in susceptibility as initial average susceptibility remains the same regardless of heterogeneity (Hébert-Dufresne et al., 2020; Shaw and Kennedy, 2021). *β*_*A*_ and *β*_*B*_ are the transmission rates for types *A* and *B* respectively and were calculated as *β*_*A*_ = *p*_*A*_*c* and *β*_*B*_ = *p*_*B*_*c* where *c* is the contact rate. Note that *c* was calculated from *R*_0,*d*_. *γ* is the recovery rate and was kept constant between the types of individuals at an assumed “true” value.

For the continuous case, we implemented the following system of ordinary differential equations derived in Elderd et al. (2008):

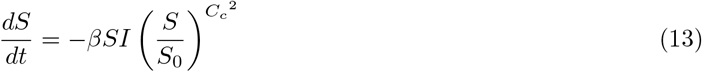

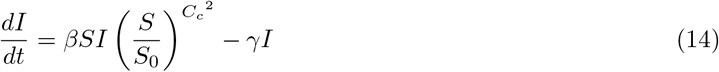

*S* is the number of susceptible individuals where *S*_0_ is the number of susceptible individuals at the beginning of the simulation, and *I* is the number of infected individuals. At the start of each simulation, we set the basic reproduction number 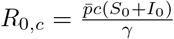 at an assumed “true” value where 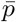 is the average probability of being infected, *c* is the contact rate, *S*_0_ + *I*_0_ is the population size, and *γ* is the recovery rate. 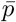is computed from the sampled parameters as 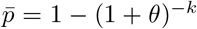, *c* was calculated from *R*_0,*c*_, and *γ* was fixed at an assumed “true” value. *β* is the transmission rate and was calculated as 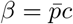.

For each case, we randomly sampled 1, 000 parameter sets from the posterior distribution to run SIR model simulations, and we compared this to the dynamics generated by the “true” parameter set used to generate our contact tracing data. Using these simulations, we determined 95% central credible intervals for the SIR dynamics for each model by finding the 2.5% and 97.5% percentiles of the 1,000 simulated dynamics at each time point over the epidemic. For our SIR simulations, we set *R*_0,*d*_ = *R*_0,*c*_ = 3, *S*_0_ = 20, 000, *I*_0_ = 10, and *γ* = 0.1.

### 2.2. Results

#### 2.2.1. Detection of heterogeneity in susceptibility

Figures 2 and 3 illustrate that the sample size, level of heterogeneity, and fraction of individuals infected affect our power to detect heterogeneity in susceptibility. This is because these factors ultimately affect the likelihoods used to test for heterogeneity in terms of the difference between the probabilities of infection for naive and focal individuals (*p*_*n*_ and *p*_*f*_) and the variability in the likelihood ratio test statistic (Supplementary information S3). More precisely, these figures show that as the number of focal individuals *F* increases from 50 to 200, there is greater power to detect lower levels of heterogeneity (lower values of *C*_*d*_, *C*_*c*_). This additionally allows for greater power across a wider range of *E*_*d*_ and *E*_*c*_. This was to be expected because higher sample sizes, particularly of the previously exposed, focal individuals, decreases variability in our estimates of *p*_*n*_, *p*_*f*_, and 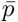. Notably, changing the total number of hosts in each contact network *N* had very little effect on our results (Supplementary information S2).

**Figure 2:**
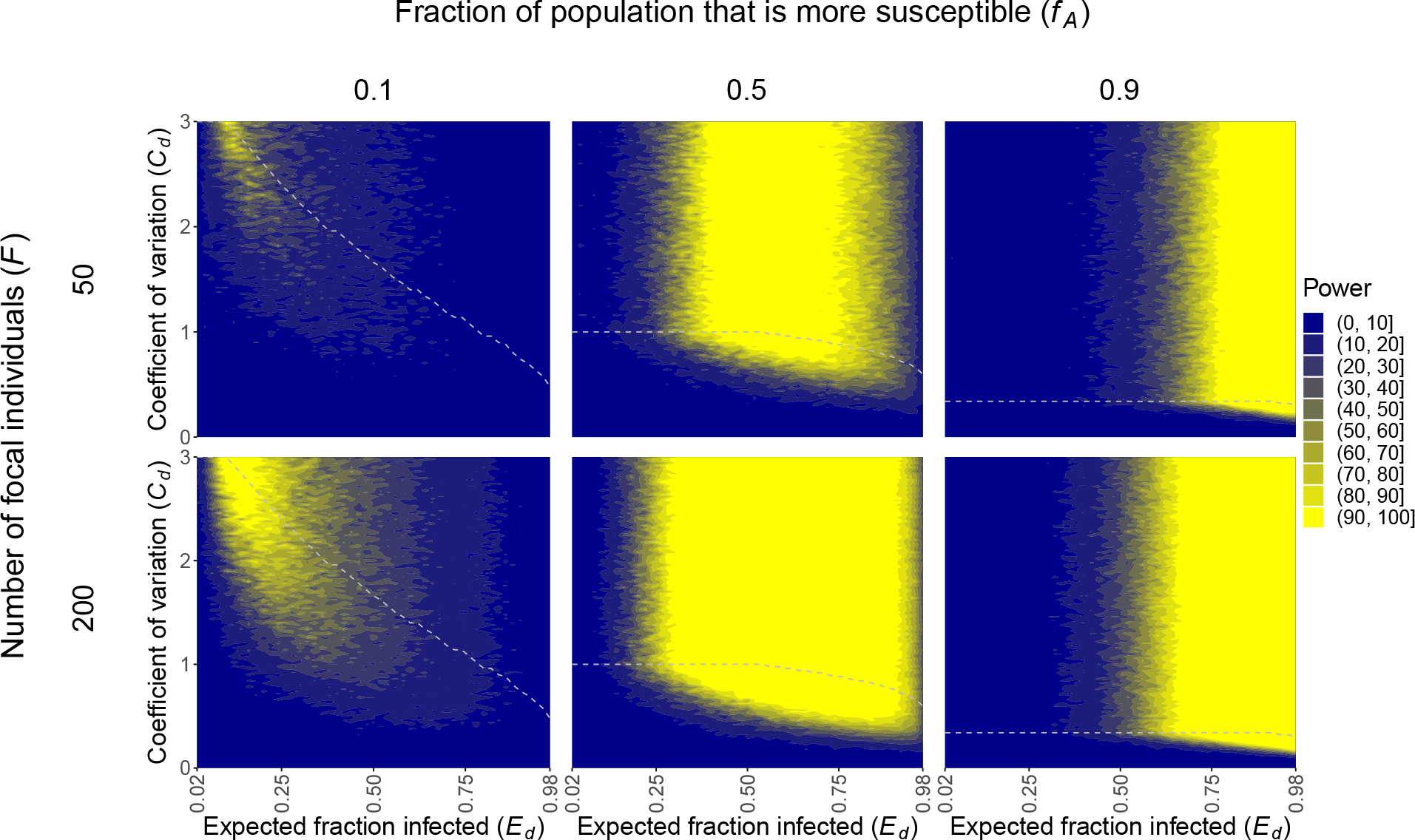
Increased heterogeneity in susceptibility (larger *C*_*d*_ and *f*_*A*_ → 0.5), intermediate fractions of individuals infected (intermediate *E*_*d*_), and increased sample sizes (larger *F*) enhance our power to detect heterogeneity in susceptibility in the discrete case. The plots show the power to detect heterogeneity in susceptibility in the discrete case across different numbers of focal individuals *F* and fraction of the population that is type *A* and more susceptible *f*_*A*_. The areas above the gray dashed lines represent parameter space that gives computationally indistinguishable probabilities of infection *p*_*A*_ and *p*_*B*_, and therefore power, to the parameter combination with the same *E*_*d*_ and highest *C*_*d*_ below the line. This occurs because risks of infection can be changed to increase *C*_*d*_ without bound, whereas probabilities are bounded. *N* = 5.

**Figure 3:**
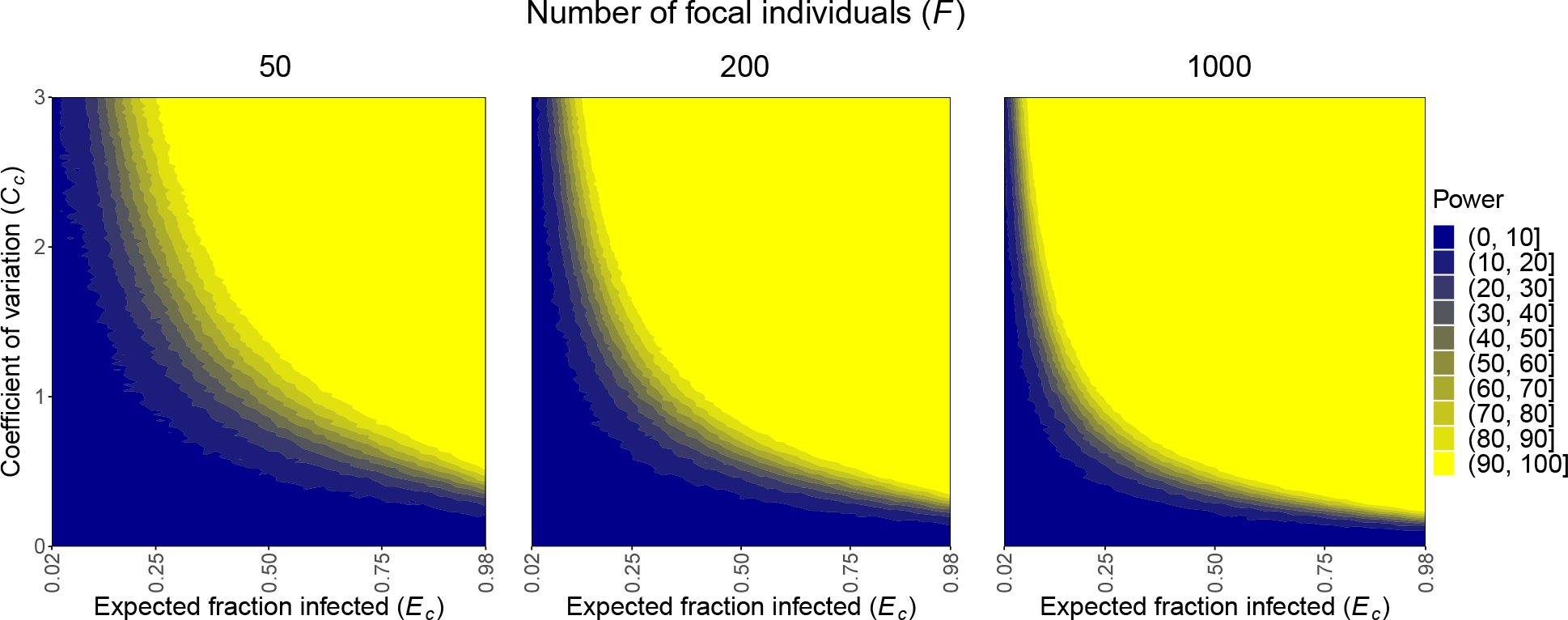
Increased heterogeneity in susceptibility (larger *C*_*c*_), greater fractions of individuals infected (larger *E*_*c*_), and increased sample sizes (larger *F*) enhance our power to detect heterogeneity in susceptibility in the continuous case. The plots show the power to detect heterogeneity in susceptibility in the continuous case across different numbers of focal individuals *F*. *N* = 5.

The level of heterogeneity in susceptibility present is described by the coefficient of variation of the risk distribution *C*_*d*_ or *C*_*c*_. As *C*_*d*_ and *C*_*c*_ increase, there is more power to detect heterogeneity in susceptibility as there is more heterogeneity in the population. In the discrete case, for a given *C*_*d*_, there is also more power to detect heterogeneity as *f*_*A*_ approaches 0.5. This is because as *f*_*A*_ approaches 0.5, the population is more evenly split between the two types of individuals, allowing for a greater difference between *p*_*A*_ and *p*_*B*_ and, therefore, *p*_*n*_ and *p*_*f*_ .

Lastly, the impact of the expected fraction of naive individuals infected (*E*_*d*_, *E*_*c*_) on power differs between the two underlying models. There is greater power to detect heterogeneity when an intermediate fraction of individuals is infected in the discrete case and when a greater fraction of individuals is infected in the continuous case. In the discrete case, *E*_*d*_ is determined by *p*_*A*_, *p*_*B*_, and *f*_*A*_ as per equation 6. The only way to have a large fraction of individuals infected is if both *p*_*A*_ and *p*_*B*_ are large. Hence, when *E*_*d*_ is high, *p*_*A*_ and *p*_*B*_ must both be close to 1. For similar reasons, when *E*_*d*_ is low, *p*_*A*_ and *p*_*B*_ must both be close to 0. Even though the risks *r*_*A*_ and *r*_*B*_ associated with these values may have varying levels of heterogeneity, the individuals themselves will have very similar infection outcomes, making it difficult to detect heterogeneity in susceptibility. Therefore, heterogeneity in susceptibility is better detected when an intermediate fraction of individuals is infected in the discrete case. In contrast, power increases in the continuous case with greater fractions of individuals infected (larger values of *E*_*c*_). This is because there is more selection for who is infected as more individuals are infected, so the average population susceptibility will decrease more drastically, making it easier to detect heterogeneity in susceptibility.

#### 2.2.2. Quantification of heterogeneity in susceptibility

We then explored the method’s ability to estimate model parameters as well as predict the associated SIR dynamics. We perform this analysis for a particular parameter combination that leads to *C*_*d*_ = *C*_*c*_ = 1.3 and *E*_*d*_ = *E*_*c*_ = 0.25. These values were chosen because they represent a biologically realistic scenario based on previous literature (Dwyer et al., 1997, 2000; De Serres et al., 2000; Rieder, 2003; Smith et al., 2005; Taylor et al., 2007; Ben-Ami et al., 2008; Elderd et al., 2008; Lessler et al., 2009; Ben-Ami et al., 2010; Pessoa et al., 2014; Ajelli et al., 2015; Langwig et al., 2017; King et al., 2018; Gomes et al., 2019; Corder et al., 2020; Koh et al., 2020; Gomes et al., 2022). In the discrete case, we used *C*_*d*_ and *E*_*d*_ and set *f*_*A*_ = 0.2 to calculate the true values *p*_*A*_ = 0.748 and *p*_*B*_ = 0.125. In the continuous case, we used *C*_*c*_ and *E*_*c*_ to calculate the true values *k* = 0.592 and *θ* = 0.626.

We determined our 95% CIs for parameter estimation of the underlying parameters with *F* = 1000 and *N* = 5 to be those shown in Table 1. Note that the true values for *p*_*A*_, *p*_*B*_, and *f*_*A*_ as well as for *k* and *θ* are captured by these intervals. Admittedly, these parameter estimates are somewhat broad. Upon further investigation, we found the broad intervals to be due to high correlation in our parameter estimates, indicating low identifiability (Fig 4, 5). However, acceptable estimates do not span the entire ranges of the parameters and encapsulate the true parameters, so there is some information about their values in the data. As we will discuss, this partial identifiability does not hinder us from making precise predictions about the impact of the heterogeneity in susceptibility on the disease dynamics.

**Figure 4:**
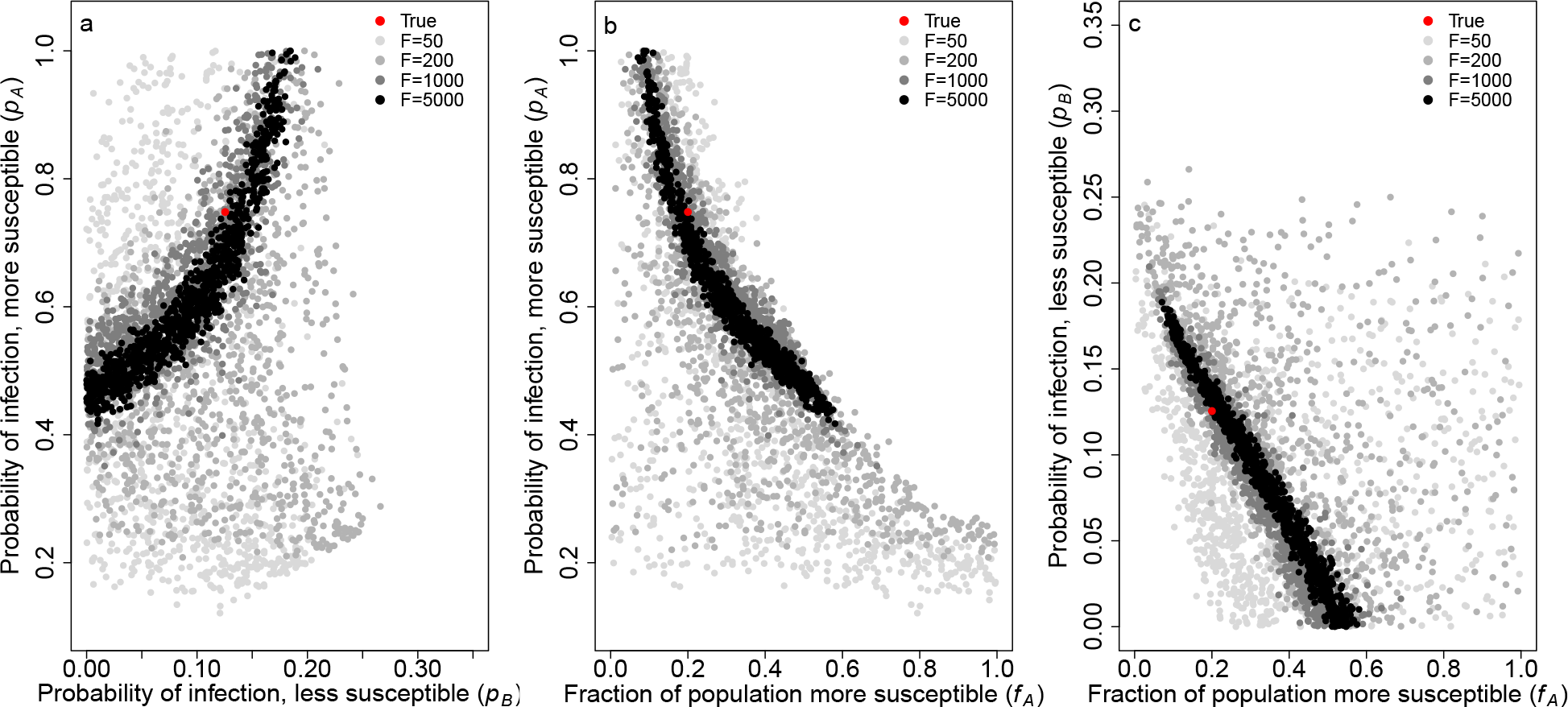
Parameter estimates for *p*_*A*_, *p*_*B*_, and *f*_*A*_ in the discrete case capture the true values and are highly correlated. The plots show the correlation in the parameter estimates for a) *p*_*A*_ vs. *p*_*B*_, b) *p*_*A*_ vs. *f*_*A*_, and c) *p*_*B*_ vs. *f*_*A*_ with different numbers of focal individuals *F*. These are the parameters that determine the distribution of individuals’ susceptibilities in the discrete case. The red dot represents the true parameters used to generate our simulated data, and the gray dots depict 1, 000 parameter sets from our posterior distribution for *F* = 50 (light gray), 200 (medium gray), 1000 (dark gray), and 5000 (black). *p*_*A*_ = 0.748, *p*_*B*_ = 0.125, *f*_*A*_ = 0.2, and *N* = 5.

**Figure 5:**
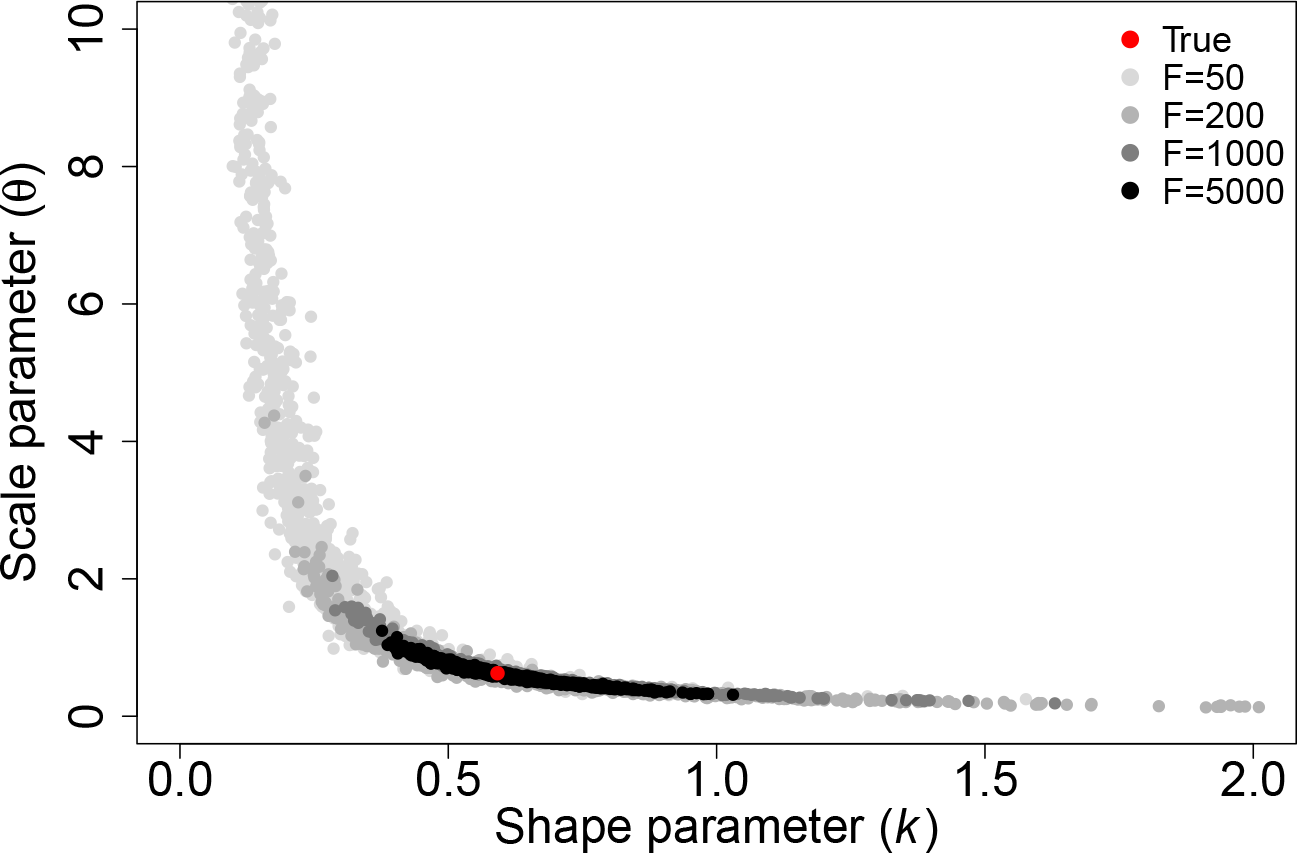
Parameter estimates for *k* and *θ* in the continuous case capture the true values and are highly correlated. This plot shows the correlation in the parameter estimates for *k* and *θ* that determine the gamma distribution of individuals’ susceptibilities in the continuous case with different numbers of focal individuals *F*. The red dot represents the true parameters used to generate our simulated data, and the gray dots depict 1, 000 parameter sets from our posterior distribution for *F* = 50 (light gray), 200 (medium gray), 1000 (dark gray), and 5000 (black). *k* = 0.592, *θ* = 0.626, and *N* = 5.

Using equations 5, 6, 8, and 9, we calculated and plotted the posterior distributions for *C*_*d*_ and *E*_*d*_ and *C*_*c*_ and *E*_*c*_ (Fig 6). With *F* = 1000 and *N* = 5, we determined the 95% CIs to be those shown in Table 1, which capture the true values. In the discrete case, the range of potential estimates for *C*_*d*_ is somewhat broad, but there is a strong ability to accurately and precisely estimate *E*_*d*_. However, in the continuous case, there is a strong ability to accurately and precisely estimate both *C*_*c*_ and *E*_*c*_. With increasing values of *F* from 50 to 5000, estimates for *C*_*c*_ and *E*_*c*_ become more precise.

**Figure 6:**
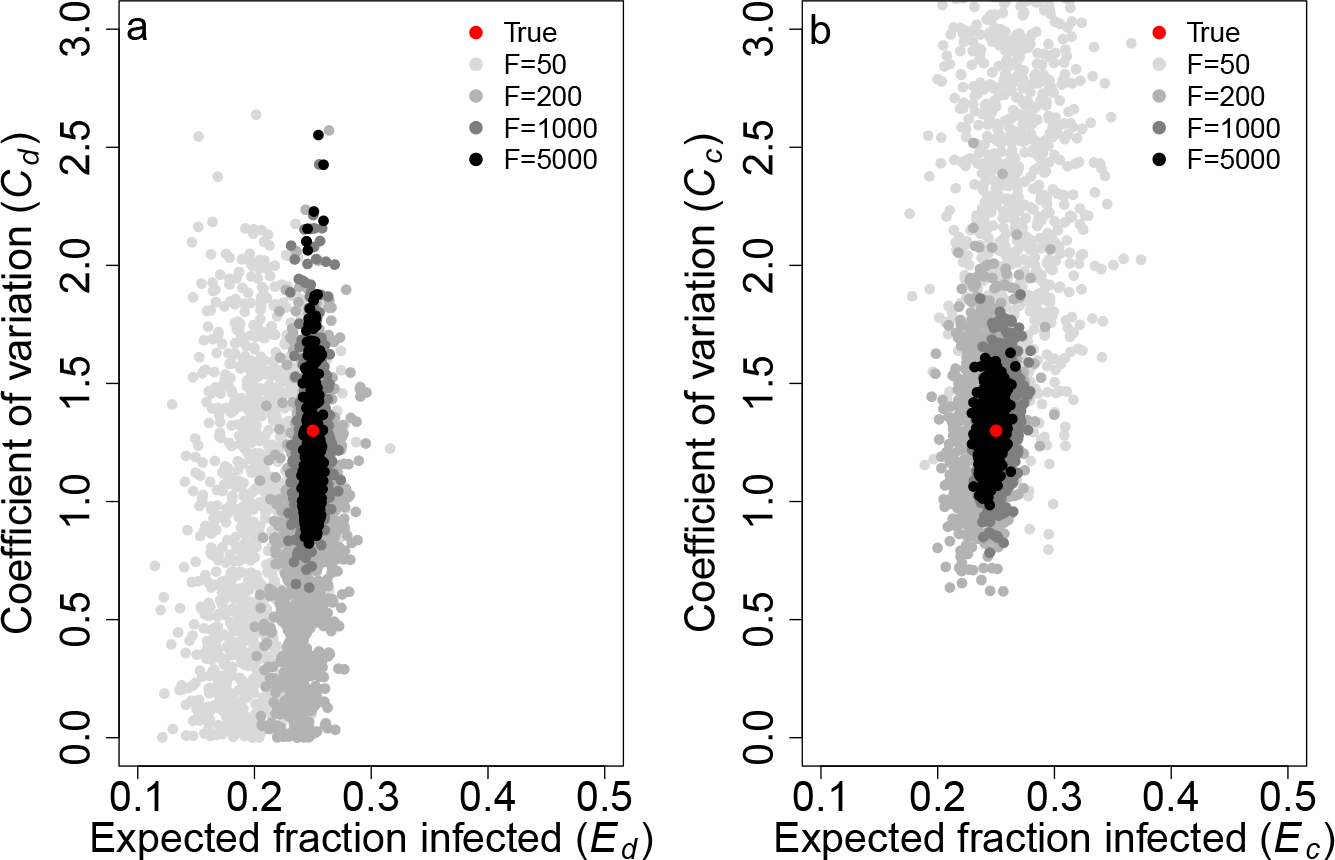
Parameter estimates for the coefficient of variation (*C*_*d*_, *C*_*c*_) and expected fraction of naive individuals infected (*E*_*d*_, *E*_*c*_) capture the true values and become more precise with increasing numbers of focal individuals *F*. The plots show the parameter estimates for *C* and *E* with different numbers of focal individuals *F* in a) the discrete case and b) the continuous case. The red dot represents the true parameters used to generate our simulated data, and the gray dots depict 1, 000 parameter sets from our posterior distribution for *F* = 50 (light gray), 200 (medium gray), 1000 (dark gray), and 5000 (black). *C*_*d*_ = *C*_*c*_ = 1.3, *E*_*d*_ = *E*_*c*_ = 0.25, *f*_*A*_ = 0.2, and *N* = 5.

We then investigated the SIR dynamics for these parameter sets with different sample sizes (*F* and *N*). We also investigated the dynamics with different error tolerances allowed for ABC in the discrete case. For both underlying models, with *N* = 5 and *F* = 50, 200, 1000, or 5000, the true dynamics are captured by the 95% CIs (Fig 7). Additionally, for *F >* 200 in the discrete case and for all *F* in the continuous case, the estimated disease dynamics do not overlap those where there is assumed to be no heterogeneity in susceptibility. Hence, despite low identifiability in the parameter estimates, we are able to use this method to make accurate and precise predictions about the effect of heterogeneity in susceptibility on disease dynamics. This is because there is interdependence among the parameters (Figs 4, 5), and so, while individual parameters may be only partially identifiable, combinations of them can be precisely estimated, leading to relatively precise estimates of the level of heterogeneity in susceptibility *C* and the fraction of naive individuals infected *E*.

**Figure 7:**
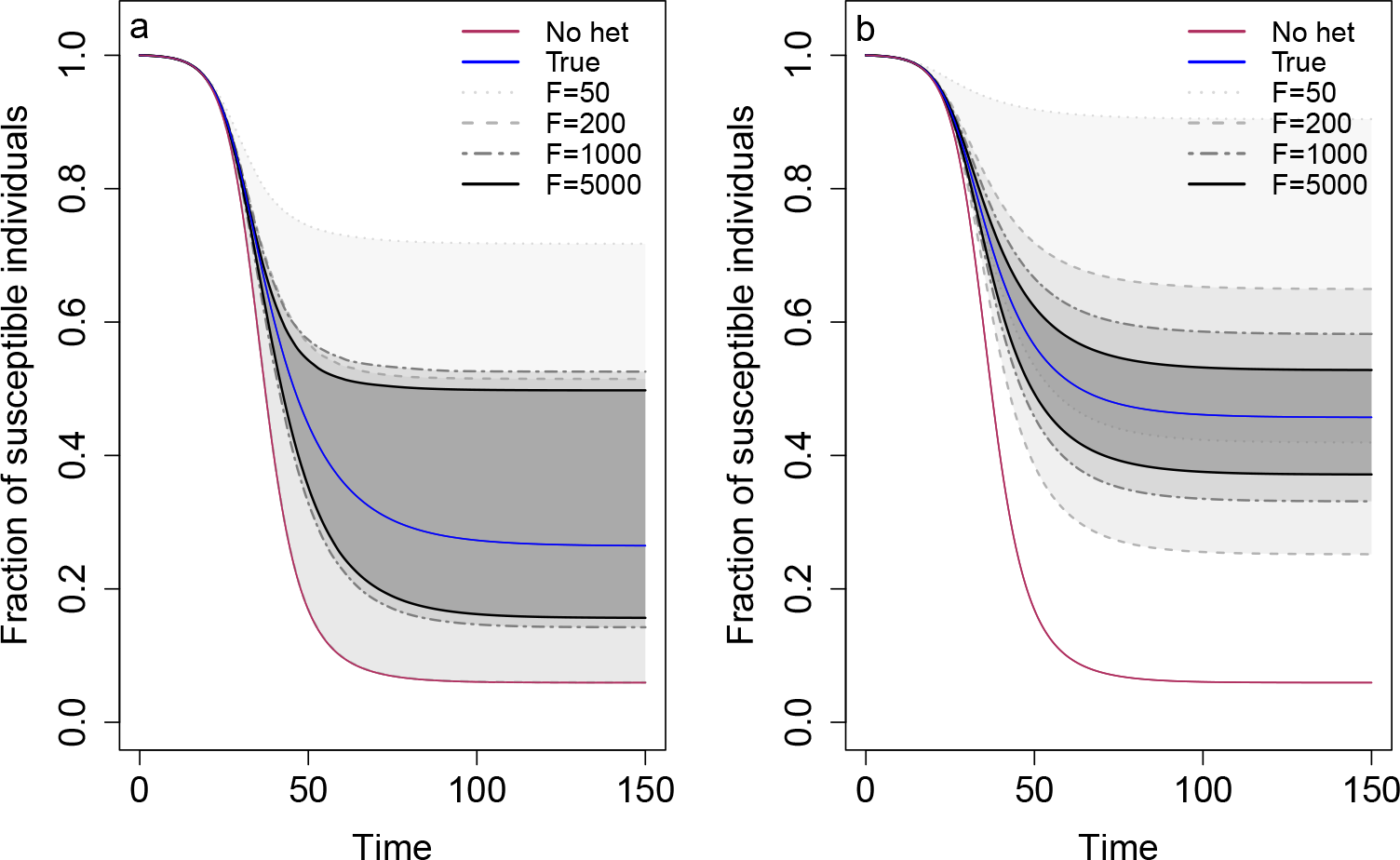
Predicted SIR dynamics capture the true dynamics and the 95% CIs narrow as the number of focal individuals *F* increases. The plots show the predicted SIR dynamics in a) the discrete case and b) the continuous case with different numbers of focal individuals *F*. Specifically, the fraction of susceptible individuals 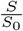 is shown over the course of an epidemic. Shaded regions represent 95% CIs determined from 1,000 posterior samples for *F* = 50 (light gray), 200 (medium gray), 1000 (dark gray), and 5000 (black). The blue line shows the true dynamics for the parameters used to generate the contact tracing data, and the red line shows the corresponding dynamics if there is homogeneity in susceptibility. *C*_*d*_ = *C*_*c*_ = 1.3, *E*_*d*_ = *E*_*c*_ = 0.25, *f*_*A*_ = 0.2, and *N* = 5.

We found the continuous case provided more accurate and precise predictions of disease dynamics than the discrete case, but the 95% CIs narrowed with higher sample sizes in both cases (Fig 7). In the discrete case, as *F* increased, there was a limit to how narrow the 95% CIs became. *F >* 1000 did not substantially improve the predicted dynamics relative to those for *F* = 1000. Likewise, the number of non-focal individuals had relatively little impact on our predicted dynamics, yielding nearly identical results for *N* = 5 and *N* = 100 (Supplementary information S2). In the continuous case, as *F* increased, the 95% CIs narrowed and converged around the true dynamics. With *N* = 5 versus *N* = 100, there was not a substantial difference in the 95% CIs (Supplementary information S2).

To assess the accuracy of our ABC method for parameter estimation in the discrete case, we examined the SIR dynamics with different error tolerances of 10%, 1%, or 0%. We did so with *N* = 5 and *F* = 200 and 1000. Changing the error tolerance did not substantially impact the precision of the 95% CIs in any of the cases explored (Supplementary information S4).

We also attempted to predict disease dynamics with the wrong underlying model of individuals’ risks as it may be unknown which model is correct in a real system. To do so, we generated data under the discrete case then predicted SIR dynamics assuming the continuous case and vice versa. Notably, the 95% CIs from the incorrectly assumed underlying models did not capture the true dynamics, meaning that caution should be taken in ensuring that an accurate model of heterogeneity is assumed before trusting the precise disease dynamics that would be expected to arise from a given set of parameter estimates (Supplementary information S5). Nevertheless, we stress that the ability to detect the presence of heterogeneity is independent of the underlying model and will not be affected by an incorrect model.

## 3. Discussion

As we saw play out during the COVID-19 pandemic, early epidemiological model predictions of disease dynamics can be crucial in informing public health policy. There are numerous imperfect assumptions made by standard SIR models, and a great deal of work has been aimed at trying to improve such models. Heterogeneity in susceptibility, differences between hosts in their likelihood of becoming infected given contact, can be critically important to disease dynamics (Dwyer et al., 1997; Gomes et al., 2014; Langwig et al., 2017; Gomes et al., 2022). However, current methods to estimate this heterogeneity rely on data that is collected late in an epidemic or is unable to be collected due to ethical or logistical constraints. Here we have developed a method to detect and estimate heterogeneity using contact tracing data which, in theory, could allow epidemiologists to incorporate the effects of heterogeneity in susceptibility into their models even before the effects of such heterogeneity are observable at the population scale. Using a simulation-based approach, we found that contact tracing data alone has enough information to be used to detect and quantify heterogeneity in susceptibility. For our method, power to detect heterogeneity increases with larger sample sizes and greater heterogeneity present as well as intermediate fractions infected in the discrete case (*E*_*d*_) and high fractions infected in the continuous case (*E*_*c*_).

Few studies have estimated heterogeneity in susceptibility in any infectious disease systems. Performing a standard literature search, we were able to find 46 estimates of heterogeneity in susceptibility from only 9 unique systems (Dwyer et al., 1997, 2000; Smith et al., 2005; Ben-Ami et al., 2008; Elderd et al., 2008; Ben-Ami et al., 2010; Pessoa et al., 2014; Langwig et al., 2017; King et al., 2018; Gomes et al., 2019; Corder et al., 2020; Gomes et al., 2022) with only 6 of those estimates pertaining to 4 human disease systems. While this list may not be entirely exhaustive, our method may be useful for expanding the set of systems for which heterogeneity in susceptibility can be detected and estimated. To determine whether our method is sufficiently powered, we need to know whether the values of the expected fraction infected *E* and the coefficient of variation of risk *C* are in a parameter space where our method would likely be suitable. Of the estimates for *C* that we found in the literature, 42 (91%) of them were greater than 0.5 and 21 (46%) were greater than or equal to 1.5. With 200 focal individuals (*F* = 200), *f*_*A*_ = 0.5, and *C* = 1.5, we have at least 80% power to detect heterogeneity in susceptibility when *E*_*d*_ is between 0.28 and 0.92 or when *E*_*c*_ is between 0.26 and 0.98. With *F* = 1000 and *C* = 1.5, we have at least 80% power when *E*_*d*_ is between 0.18 and 0.98 or when *E*_*c*_ is between 0.14 and 0.98 (Figs 2, 3). In studies examining contact tracing data, we found secondary attack rates, which provide conservative estimates of *E*, to often be around 0.2 and sometimes as high as 0.733 (De Serres et al., 2000; Rieder, 2003; Taylor et al., 2007; Lessler et al., 2009; Ajelli et al., 2015; Koh et al., 2020). Our method should therefore be sufficiently powered for many systems.

The precision in our prediction of SIR dynamics is also affected by the nature of the heterogeneity in susceptibility. Our estimates of how heterogeneity affects disease dynamics are less precise when there are discrete differences in risk between hosts, as opposed to continuous variation in risk (Fig 7). This is because, in addition to *C*_*d*_ and *E*_*d*_, the fraction of the initial population that is the more susceptible type of individual, *f*_*A*_, is critical for determining the trajectory of the epidemic. With the same *C*_*d*_ and *E*_*d*_, the final epidemic size can differ depending on *f*_*A*_ (Supplementary information S6). Hence, the need to estimate the additional parameter *f*_*A*_ in the discrete case with the same data results in wider 95% CIs. However, we can generate narrow 95% CIs and more precise parameter estimates in the discrete case if there is prior knowledge of the parameters *p*_*A*_, *p*_*B*_, or *f*_*A*_ (Supplementary information S7).

We found that using the correct underlying model is additionally important for accurately predicting disease dynamics, but not for the detection of heterogeneity in the first place. The underlying model used for parameter estimation should therefore be carefully chosen to reflect prior understanding of the potential drivers of heterogeneity in susceptibility in the system. The process for initial detection of heterogeneity in susceptibility is the same regardless of the underlying model (Eqs. 1, 2). Therefore, we can reliably detect heterogeneity in susceptibility without knowledge of the distribution of individuals’ risks.

One strength of our method is that it allows for estimation of heterogeneity in susceptibility in real time, early in an epidemic with no data other than contact tracing data. Admittedly, the use of this data in real time will depend on the speed with which the necessary data can be collected and communicated, but existing methods to quantify heterogeneity are not adequate for real time usage even with immediate access to the data. Ben-Ami et al. (2010) and Langwig et al. (2017) used experimental dose-response curves to estimate heterogeneity in susceptibility, and Dwyer et al. (1997) used a combination of laboratory dose-response experiments, field transmission experiments, and models fit to mortality data to investigate heterogeneity. Although these experimental methods can provide good estimates of heterogeneity in susceptibility, they are not feasible for application in real time or for human epidemics in general due to time constraints and ethical concerns. Gomes et al. (2019) compared disease incidence across municipalities in several countries to quantify heterogeneity for tuberculosis. This was done by ordering the municipalities by incidence rate and plotting the percentage of cumulative tuberculosis cases versus cumulative population to construct Lorenz curves and thereby fit susceptibility risk distributions. This method, however, requires a considerable amount of data with ten or more years of data used in this study. Smith et al. (2005) and Corder et al. (2020) used malaria morbidity data to fit models of malaria and estimate heterogeneity. This method cannot be used until later in an epidemic when sufficient data is collected to fit curves. Gomes et al. (2022) also used curve fitting with mortality data to estimate heterogeneity in susceptibility for COVID-19. They were able to estimate heterogeneity in real time once at least four months of data were available. While our method is in principle able to estimate heterogeneity in a similar time frame provided robust contact tracing, we also note that their method is heavily dependent on the underlying model and assumptions, and the authors advise not to trust the precision of their estimates. In addition, Gomes et al. were unable to disentangle heterogeneity in contact rate from heterogeneity in underlying susceptibility. Our method estimates heterogeneity in underlying susceptibility, and the remaining heterogeneity in contact rate can be determined from the contact network data. Anderson et al. (2023) used household study data to estimate heterogeneity in susceptibility. While this method is suitable for use in real time, and can be applied to human infectious diseases, the method notably is designed to estimate heterogeneity within households, which is not the same as the population-level heterogeneity that drives population-level disease dynamics.

Our method is unable to precisely estimate the individual parameters that define the risk distributions (i.e. *p*_*A*_, *p*_*B*_, *f*_*A*_ in the discrete case and *k, θ* in the continuous case), but our method is able to reliably predict disease dynamics. This seeming paradox arises because the disease dynamics depend on combinations of parameters rather than individual parameters. Notably, our method is substantially better at estimating the composite parameters describing the coefficient of variation of risk *C* and the expected fraction of naive individuals infected *E*. Nevertheless, our method does require a substantial amount of data (200 individuals showing up in contact networks for a second time). This requirement could be mitigated by pooling contact network data from multiple locations in order to more quickly collect sufficient data. It may also be possible to combine our method with another, like that of Gomes et al. (2022), to reduce the data required by either method.

There are additionally several considerations to address with regard to working with contact tracing data. Perhaps most prominently, contact tracing data tend to be messy and imperfect. Our method as described above assumes perfect data. However, our method can be readily modified to account for imperfect data. We can imagine multiple ways in which contact tracing data may be imperfect. Some important considerations are that: a) individuals may be mislabeled as uninfected when they are infected (false negatives), b) individuals may be mislabeled as infected when they are uninfected (false positives), and c) individuals may be missing from the contact networks despite being contacts (missing contacts). If there are false negatives, our method may overestimate the level of heterogeneity because our estimate of *p*_*f*_ may be biased lower. This is because, assuming infection confers at least partial immunity, focal individuals that were actually infected previously (i.e. false negatives) will be less likely to be infected than focal individuals that were true negatives. To counteract this issue, we developed a version of the method that corrects for false negatives by adjusting the likelihood calculations for both detecting and estimating heterogeneity. For estimating parameters and predicting disease dynamics, adjusting the method to correct for false negatives fixes the issue (Supplementary information S8). For detecting heterogeneity in susceptibility, adjusting the likelihood calculation corrects for the impact of false negatives except when the expected fraction infected *E*_*d*_ is very close to 1. We do not think this will be a major issue as *E*_*d*_ is typically less than 0.5 (De Serres et al., 2000; Rieder, 2003; Taylor et al., 2007; Lessler et al., 2009; Ajelli et al., 2015; Koh et al., 2020). If there are false positives, our method may underestimate the level of heterogeneity because our estimate of *p*_*f*_ may be biased higher. This is because a high false positive rate will have a larger impact on making individuals with a low susceptibility appear infected than those with a high susceptibility that are more likely to be true positives. Hence, focal individuals, which are on average less susceptible, and naive individuals will appear to have more similar infection probabilities. However, false positive rates are often small, close to 1-2% (Yang and Rothman, 2004; Cohen et al., 2020), so this issue is not a huge concern for our method unless false positive rates are known to be unusually large. If there are many missing contacts, our method could underestimate the level of heterogeneity because our estimate of *p*_*n*_ may be biased lower. This is because individuals that we believe to be naive but were previously exposed in a first contact network may be less likely to be infected than true naive individuals. These missed individuals may have gained immunity through infection or may be on average less susceptible through the infection selection process. However, there is a low chance of a missed individual from a first contact network showing up in a second contact network that also happens to have a focal individual early in an epidemic. So, missing individuals should have only a negligible effect on the method’s performance in these early stages. Later on in an epidemic, this source of bias will become more important to consider. While we have considered these three ways in which contact tracing data may be imperfect, it is highly likely that each set of contact tracing data will have its own set of peculiarities. Note that these peculiarities, if known, can readily be accounted for using our ABC method since any process may be used for simulation. Known imperfections in the data should therefore not bias estimates although they may still reduce power or increase required sample sizes.

Another important point is that our method assumes no forms of heterogeneity other than heterogeneity in susceptibility. One other source of heterogeneity is heterogeneity in transmission (Lloyd-Smith et al., 2005). Heterogeneity in transmission is differences between hosts in their likelihood of transmitting a pathogen once infected. If this heterogeneity arises due to variation in the number of contacts that individuals have, then heterogeneity in transmission poses no problems for our method. It would simply mean that each contact network would have a unique value for *N*. We note that this variation in contact rate is the typical mechanism through which heterogeneity in transmission is assumed to act (Lloyd-Smith et al., 2005). However, if heterogeneity in transmission arises due to differences between hosts in their likelihood of transmission given contact, our method may have less power to detect heterogeneity in susceptibility and may yield less precise or faulty conclusions about the disease dynamics (Supplementary information S9). Our method, in its existing form, is thus not suitable in these cases. This concern can be partially mitigated by performing a goodness of fit test before implementing the method to determine whether there is evidence of heterogeneity in transmission given contact (Supplementary information S9). If there is heterogeneity in transmission, then our method should not be used. A next step in developing this method will be to generalize it to allow for estimation of heterogeneity in susceptibility even when there is heterogeneity in transmission. This is, however, a non-trivial problem because if every individual has a unique force of infection, then the number of parameters to estimate grows at the same rate as the number of focal individuals.

There may additionally be heterogeneity in exposure strength among contacts within a network such that individuals experience different forces of infection. This could be due to factors like differences in exposure time or type of contact (e.g., contacts that shared a taxi, were at the same party, etc.). This added heterogeneity may reduce the power of our method to detect heterogeneity in susceptibility as different contact types may provide varying levels of information that our current method disregards. To alleviate the potential impact of this heterogeneity, it may be necessary to break apart contact networks into specific exposure events and either weigh the type of contact differently or only use equivalent contact types.

Finally, we note that exposure could change individuals’ susceptibilities. Individuals exposed in a first contact network could receive a small dose of the pathogen such that their immune system is stimulated without them becoming infected. This could decrease their susceptibility, meaning that some focal individuals have lower susceptibilities because they developed immunity, not because they were innately less susceptible (Leon and Hawley, 2017). However, this will have the same effect as heterogeneity in susceptibility of slowing down the epidemic and could even be considered a form of heterogeneity in susceptibility.

The earliest practice of tracing diseases dates back to the 1500s when doctors would track the spread of syphilis (Cohn, 2018), and the earliest known example of contact tracing dates to 1576 during a bubonic plague pandemic (Cohn, 2009). Since then, the practice of contact tracing has spread, and it is now used widely, ranging from diseases such as influenza to HIV (De Serres et al., 2000; Rieder, 2003; Taylor et al., 2007; Lessler et al., 2009; Ajelli et al., 2015; Koh et al., 2020). Recently, contact tracing data has transitioned from paper copies to electronic databases. Regardless, all of these sources of data could be used with our method provided they include focal individuals that are identifiable between contact networks, specify which individuals are infected, and have a sufficient sample size. Using our method, it should therefore, without collecting any new data, be possible to estimate heterogeneity in susceptibility, in various locations and time periods, for dozens of disease systems in which it has never been estimated previously.

## Acknowledgements

We thank the Read, McGraw, and Kennedy labs for stimulating discussions.

