## Supplementary Information for "Detecting and quantifying heterogeneity in susceptibility using contact tracing data"

### S1. Derivation of $C_d$

The coefficient of variation is defined as standard deviation divided by the mean. Hence, $C_d$ is the standard deviation of risk divided by the mean risk.

The mean risk, $\mu_r$, is given by

$$\mu_r = E(\text{risk}) = r_A f_A + r_B(1 - f_A),$$

and the standard deviation of risk, $\sigma_r$, is given by

$$\begin{aligned}\sigma_r &= \sqrt{\sigma_r{}^2} \\ &= \sqrt{E(\text{risk}^2) - E(\text{risk})^2} \\ &= \sqrt{(r_A{}^2 f_A + r_B{}^2(1 - f_A)) - (r_A f_A + r_B(1 - f_A))^2}.\end{aligned}$$

$C_d = \frac{\sigma_r}{\mu_r}$ can then be simplified to

$$C_d = \frac{(r_A - r_B)\sqrt{f_A(1 - f_A)}}{r_A f_A + r_B(1 - f_A)}.$$

### S2. Changing $N$

For both underlying models, we explored the effect of changing the number of individuals in each contact network $N$ on our power to detect heterogeneity in susceptibility and our ability to predict SIR dynamics. For detection, we followed the same method described in the main text with $N = 5$ or $N = 100$, $F = 200$, and $f_A = 0.5$ then compared the difference in power for these sample sizes. For parameter estimation and disease dynamics prediction, we followed the same method described in the main text for $N = 100$ with $C_d = C_c = 1.3$, $E_d = E_c = 0.25$, and $f_A = 0.2$. Since both $N$ and $F$ affect the sample size, we did this for $F = 200$ and $F = 5000$ to check if $N$ had a different effect with different $F$. To estimate parameters with $N = 5$, we used the same contact networks simulated for $N = 100$. This allowed us to compare differences due to $N$ while minimizing stochasticity from simulating the data. To do so, we randomly sampled the number of infected naive individuals $x_{i,s}$ in each network $i$ with $N = 5$ based on the fraction of naive individuals infected $\frac{x_{i,l}}{99}$ in $i$ with $N = 100$. Thus, $x_{i,s}$ has distribution $\text{Binom}(n = 4, p = \frac{x_{i,l}}{99})$. Since there is one focal individual in each network regardless of $N$, we used the same number of focal individuals infected (0 or 1) in $i$ for $N = 5$ as simulated for $N = 100$. After determining $x_{i,s}$ for all $i$, we ran MCMC and generated SIR dynamics as before.

We found that our power to detect heterogeneity in susceptibility and our ability to predict SIR dynamics were not substantially affected by increasing the number of individuals in each contact network, $N$, from 5 to 100 (Figs S1, S2, S3, S4, S5). In certain parameter space, $N$ may have some importance for detection, but power is overall not that sensitive to $N$ (Figs S1, S2, S3, S4). We might have expected that higher $N$, or more naive individuals, would decrease variability in our estimate of $p_n$ and allow us to more precisely estimate parameters. However, there are $F(N - 1)$ naive individuals in the second exposure round, so even when $N$ is small there is not much variation in the fraction of naive individuals infected.

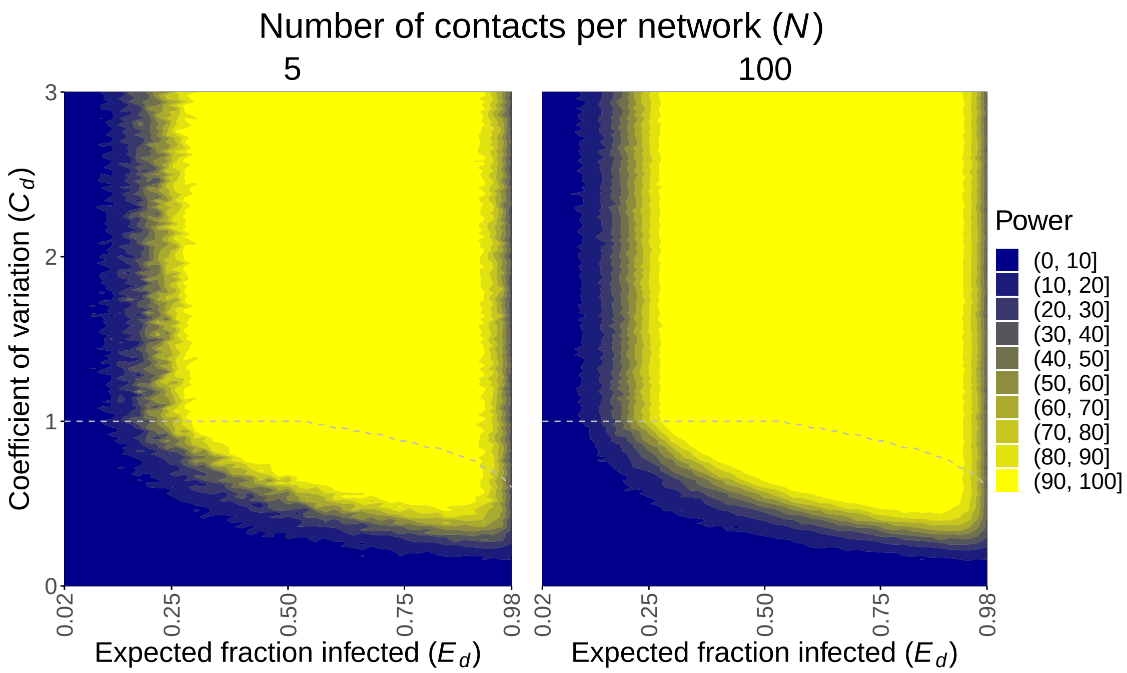


Figure S1: Power to detect heterogeneity in susceptibility in the discrete case across different numbers of individuals in a contact network $N$. The areas above the gray dashed lines represent parameter space that gives computationally indistinguishable probabilities of infection $p_A$ and $p_B$, and therefore power, to the parameter combination with the same $E_d$ and highest $C_d$ below the line. This occurs because risks of infection can be changed to increase $C_d$ without bound, whereas probabilities are bounded. $F = 200$ and $f_A = 0.5$.

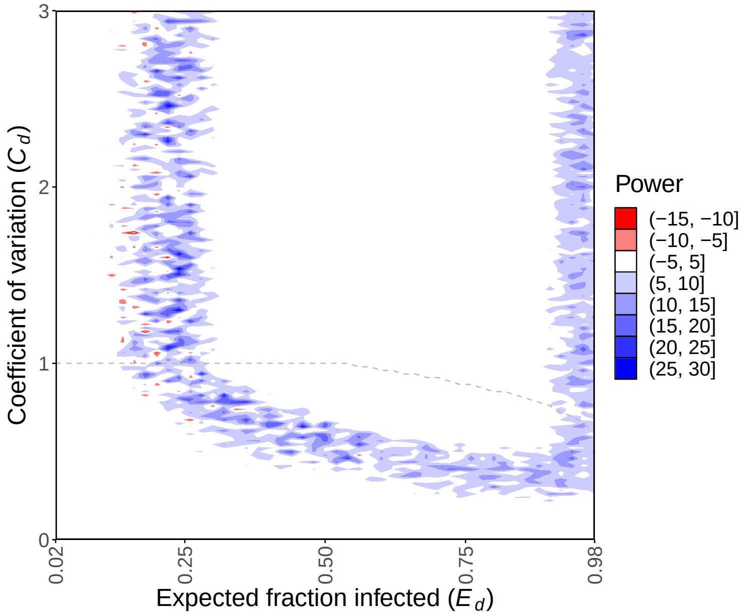


Figure S2: There is generally more power to detect heterogeneity in susceptibility in the discrete case when $N = 100$ than when $N = 5$, but the effect is relatively small and restricted to the space where heterogeneity is sometimes detectable with either value of $N$. This plot shows the difference in the power to detect heterogeneity in susceptibility between $N = 100$ and $N = 5$ for the discrete case. Positive (blue) areas mean that there is more power to detect heterogeneity in susceptibility with $N = 100$, and negative (red) areas mean there is less power with $N = 100$. $F = 200$ and $f_A = 0.5$.

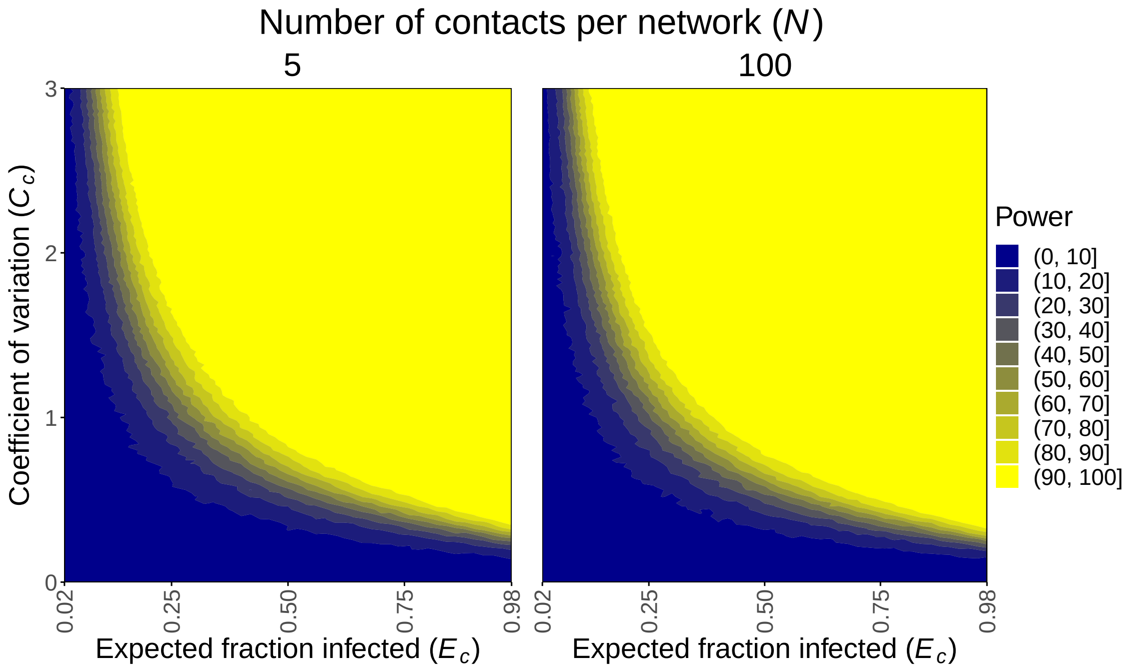


Figure S3: Power to detect heterogeneity in susceptibility in the continuous case across different numbers of individuals in a contact network $N$. $F = 200$.

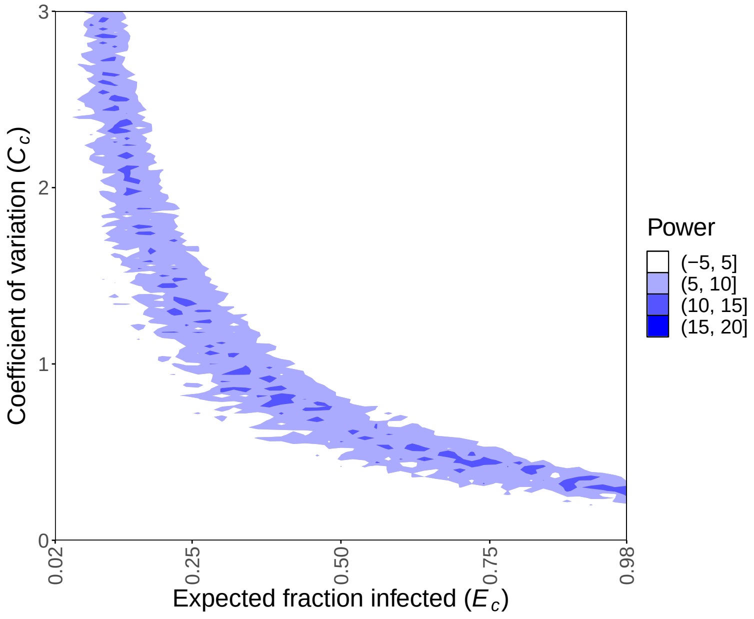


Figure S4: There is generally more power to detect heterogeneity in susceptibility in the continuous case when $N = 100$ than when $N = 5$, but the effect is relatively small and restricted to the space where heterogeneity is sometimes detectable with either value of $N$. This plot shows the difference in the power to detect heterogeneity in susceptibility between $N = 100$ and $N = 5$ for the continuous case. Positive (blue) areas mean that there is more power to detect heterogeneity in susceptibility with $N = 100$. $F = 200$.

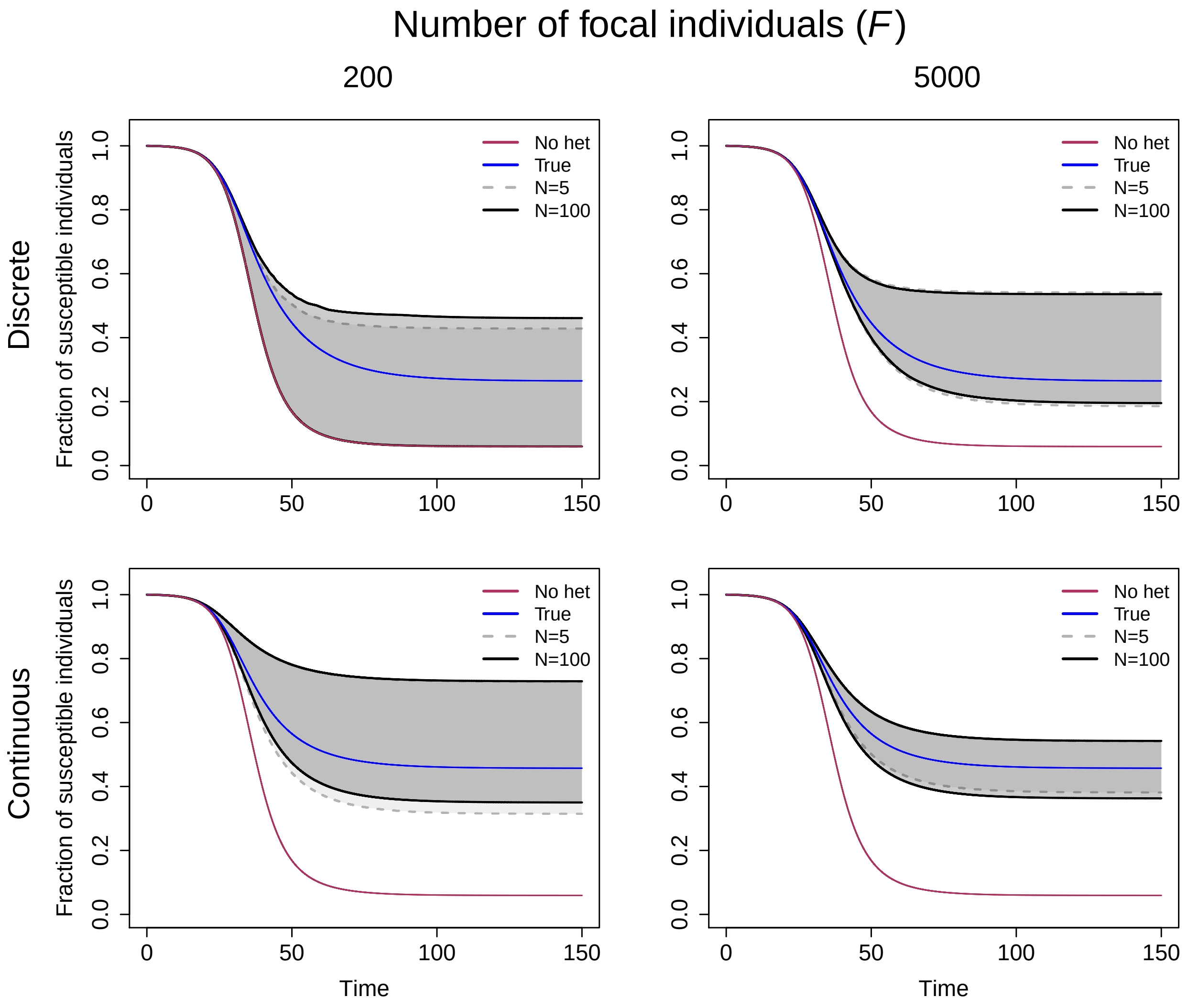


Figure S5: The number of contacts $N$ does not substantially affect our prediction of the disease dynamics. The plots show the predicted SIR dynamics with $N = 5$ versus $N = 100$ contacts per network for both the discrete and the continuous case and different numbers of focal individuals $F$. Specifically, the fraction of susceptible individuals $\frac{S}{S_0}$ is shown over the course of an epidemic. Shaded regions represent 95% CIs determined from 1,000 posterior samples for $N = 5$ (gray) and $N = 100$ (black). The blue line shows the true dynamics for the parameters used to generate the contact tracing data, and the red line shows the corresponding dynamics if there is homogeneity in susceptibility. $C_d = C_c = 1.3$, $E_d = E_c = 0.25$, and $f_A = 0.2$.

### S3. Difference between probabilities of infection for naive and focal hosts ($p_n$ and $p_f$) and variability in the likelihood ratio test statistic

We found that our power to detect heterogeneity in susceptibility increased with larger sample sizes and greater heterogeneity present as well as an intermediate expected fraction of naive individuals infected $E_d$ in the discrete case and a high expected fraction infected $E_c$ in the continuous case. We wanted to further understand how these factors affect our power in terms of the likelihoods used to test for heterogeneity and the probabilities of infection for naive, focal, and average hosts $p_n$, $p_f$, and $\bar{p}$ in them. As $p_n$ and $\bar{p}$ are typically very similar, we focused on investigating $p_n$ and $p_f$. We thought that the difference between $p_n$ and $p_f$, $\Delta p = p_n - p_f$, may be important for detecting heterogeneity. To check this, we tested the hypothesis that $p_n$ and $p_f$ are different from each other against the null hypothesis that they are the same. We did this for $p_n \in [0, 1]$ with step size 0.01 and $p_f \in [0, p_f]$ with step size 0.01. Note that although we tested each combination of $p_n$ and $p_f$ to better understand and visualize our ability to detect heterogeneity, not all combinations are possible for data with coefficient of variation of risk $C \in [0, 3]$ and expected fraction infected $E \in [0.02, 0.98]$. In figure S6, the area above the gray line is the parameter space that is possible in the discrete case with $F = 50$, $N = 5$, and $f_A = 0.5$.

For each parameter combination, we first set the probabilities of infection for naive and focal hosts $p_n$ and $p_f$ and simulated the number of naive and focal individuals infected with $F = 50$ and $N = 5$ where the number of naive individuals infected has distribution $\text{Binom}(n = F(N-1), p_n)$, and the number of focal individuals infected has distribution $\text{Binom}(n = F, p_f)$. We calculated the log-likelihood of the data the same as in the main text where $L_{\text{hom}}$ is the log-likelihood under the null hypothesis that $p_n$ and $p_f$ are the same, and $L_{\text{het}}$ is the log-likelihood under the alternative hypothesis that $p_n$ and $p_f$ are different and there is, therefore, heterogeneity in susceptibility. We then compared the log-likelihoods of the data under each hypothesis using a likelihood ratio test with one degree of freedom and significance level $\alpha = 0.05$. We ran $10,000$ simulations for each set of parameters to determine our statistical power to detect a difference between $p_n$ and $p_f$ or heterogeneity with that parameter combination.

Figure S6 shows that power increases as the difference in probability of infection $\Delta p$ increases. But, the contour lines are curved such that there is less power as $p_n, p_f \to 0.5$ and more power as $p_n, p_f \to 0$ or $p_n, p_f \to 1$. This is because there is stochasticity in which individuals become infected, and, with a binomial distribution, the highest variance is at a probability of 0.5. This stochasticity then leads to variability in the likelihood ratio test statistic, making it harder to detect a difference between $p_n$ and $p_f$. Hence, there is less power as $p_n, p_f \to 0.5$ and the variance increases. Overall, we found that our ability to detect heterogeneity in susceptibility ultimately depends on $\Delta p$ and variability in the likelihood ratio test statistic stemming from stochasticity in exposure events.

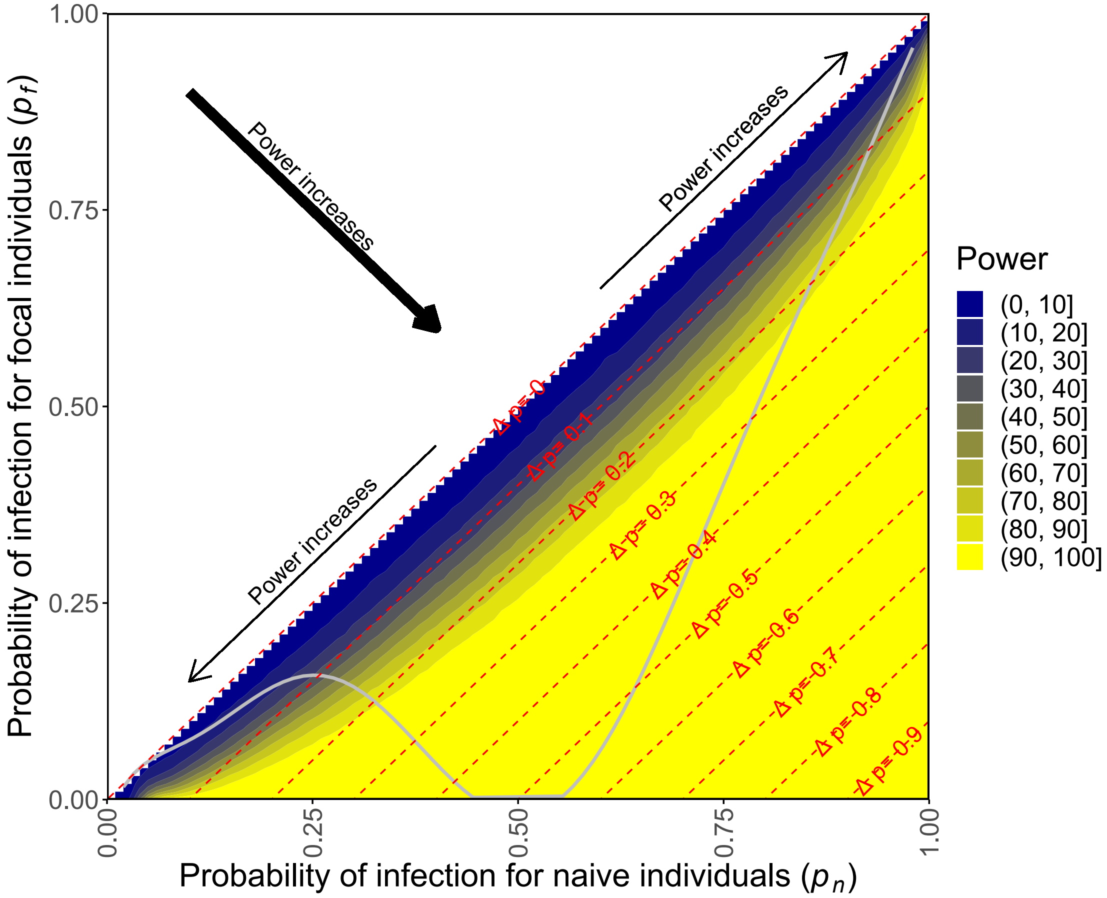


Figure S6: The effects of $\Delta p = p_n - p_f$ and variability in the likelihood ratio test statistic on the power to detect heterogeneity in susceptibility. Power increases substantially as $\Delta p$ increases, and power also increases slightly as variability in the test statistic decreases (i.e. as $p_n, p_f \to 0$ or $p_n, p_f \to 1$). The red dashed lines correspond to the values of $\Delta p$, and the arrows show how power increases with their thickness reflecting importance. The area above the gray line is the parameter space that is possible in the discrete case. All parameter space is possible in the continuous case given a large enough coefficient of variation $C_c$. $F = 50$, $f_A = 0.5$, and $N = 5$.

### S4. Changing error tolerance for Approximate Bayesian Computation (ABC) in the discrete case

To estimate parameters using our method with data that follows a discrete distribution of susceptibilities, we used ABC (Beaumont et al., 2002). This ABC method estimated the likelihood by simulating the data 100 times and examining the fraction of simulations that were close to our observed data. Notably, "close" can be defined in many ways. Here we explored the effect of our definition of close on the outcome of the model. To do so, we ran ABC where we estimated the likelihood from the fraction of simulations where the number of individuals infected was within a 10%, 1%, or 0% error tolerance of the number infected in the observed data with $F = 200$ or 1000 and $N = 5$. We then compared the resulting predicted SIR dynamics. We used two different $F$ to check if the error tolerance allowed was more important for different sample sizes. We found that changing the error tolerance did not substantially impact the precision of the 95% CIs. This can be seen in Figure S7 by comparing the CIs for an error tolerance of 10% (light gray), 1% (dark gray), and 0% (black) in each panel.

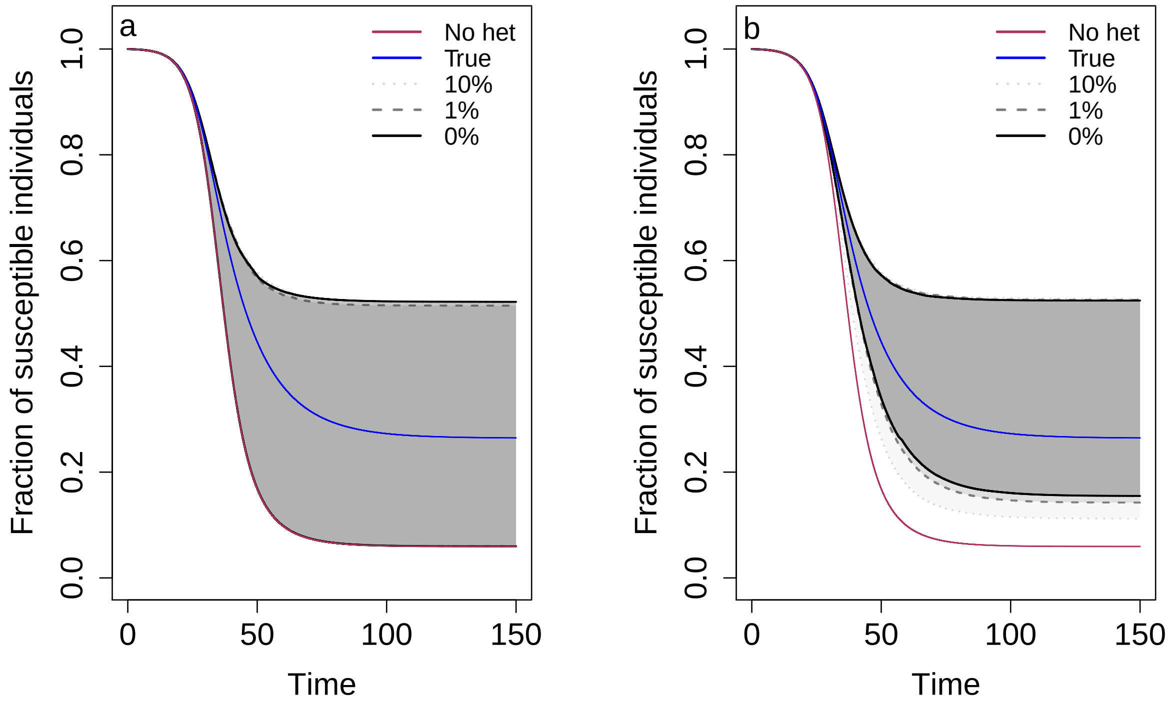


Figure S7: The error tolerance allowed for ABC has little effect on our prediction of the disease dynamics. The plots show the predicted SIR dynamics in the discrete case with different error tolerances allowed for ABC with a) $F = 200$ and b) $F = 1000$. Specifically, the fraction of susceptible individuals $\frac{S}{S_0}$ is shown over the course of an epidemic. Shaded regions represent 95% CIs determined from 1,000 posterior samples for an error tolerance of 10% (light gray), 1% (dark gray), and 0% (black). The blue line shows the true dynamics for the parameters used to generate the contact tracing data, and the red line shows the corresponding dynamics if there is homogeneity in susceptibility. $C_d = 1.3$, $E_d = 0.25$, $f_A = 0.2$, and $N = 5$.

### S5. Effect of assuming the wrong underlying model

When estimating parameters with our simulated data, we know whether individuals' risks follow a discrete or continuous distribution. However, in a real system, it may be unknown which underlying model is correct. We therefore explored the impact of assuming the wrong underlying model on our estimated parameters and predicted disease dynamics. To do so, we generated data under the discrete case then predicted SIR dynamics assuming the continuous case and vice versa. Figure S8 shows that the 95% CIs from the incorrectly assumed underlying models did not capture the true dynamics in either case. This is because the way individuals' risks are distributed in each underlying model results in fundamentally different epidemics. In the continuous case, there is a core group of highly resistant individuals that leads to a smaller epidemic size, whereas in the discrete case, when $f_A = 0.2$, the less susceptible type $B$ individuals are not highly resistant ($p_B = 0.125$), leading to a larger epidemic. However, in the discrete case, when $f_A = 0.5$, the type $B$ individuals are much more resistant ($p_B = 7.4e-05$), so the 95% CIs from the incorrectly assumed continuous case better captured the true dynamics (Fig S8). Overall, the underlying model used should be carefully chosen to reflect prior understanding of the potential drivers of heterogeneity in susceptibility in the system. Nevertheless, our process for detecting heterogeneity in susceptibility does not depend on the underlying model, so we can reliably detect heterogeneity without knowledge of the distribution of individuals' risks.

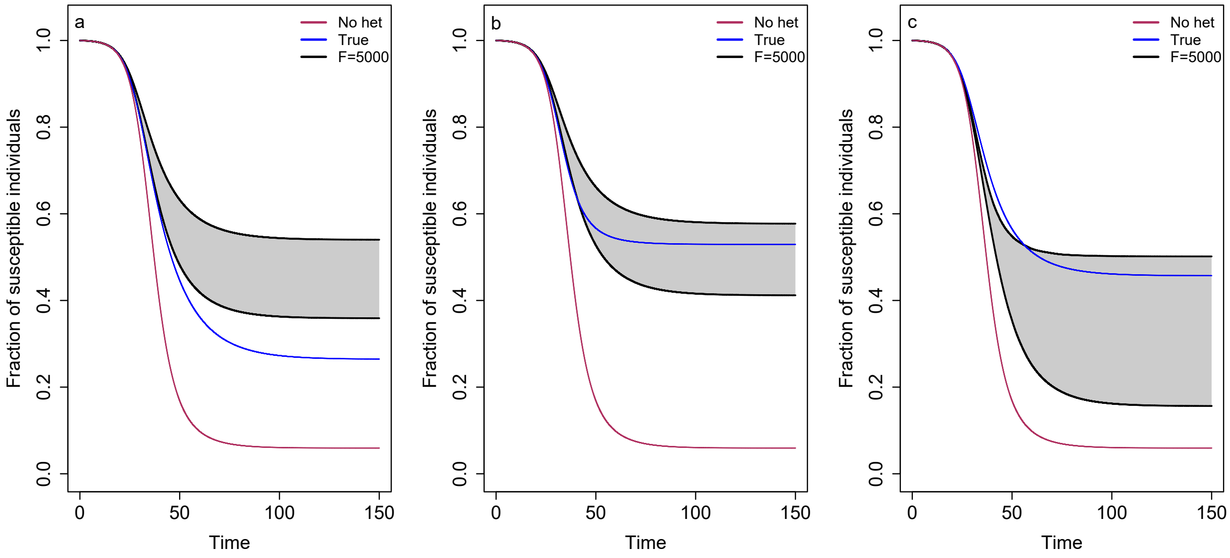


Figure S8: Assuming the wrong underlying model for estimation results in incorrect predictions of the disease dynamics, but this effect is lessened as $f_A \to 0.5$ when the discrete case is the correct model. The plots show the predicted SIR dynamics with the wrong underlying model assumed for a) the discrete case with the continuous case incorrectly assumed and $f_A = 0.2$, b) the discrete case with the continuous case incorrectly assumed and $f_A = 0.5$, and c) the continuous case with the discrete case incorrectly assumed. Specifically, the fraction of susceptible individuals $\frac{S}{S_0}$ is shown over the course of an epidemic. Shaded regions represent 95% CIs determined from 1,000 posterior samples for $F = 5000$. The blue line shows the true dynamics for the parameters used to generate the contact tracing data, and the red line shows the corresponding dynamics if there is homogeneity in susceptibility. In each plot, the data was simulated according to the true underlying model, but parameters were estimated according to the wrong underlying model. $C_d = C_c = 1.3$, $E_d = E_c = 0.25$, and $N = 5$.

### S6. Final epidemic size in the discrete case with different fractions of the population that are more susceptible ($f_A$)

When testing our method with different parameter combinations for the discrete case, we noticed that, with the same expected fraction infected $E_d$, we generated different powers to detect heterogeneity in susceptibility and different SIR dynamics by changing the fraction of individuals that was more susceptible $f_A$, even if the coefficient of variation of risk $C_d$ was kept constant. To better understand the effect of $f_A$ on disease dynamics in relation to $C_d$, we investigated the final epidemic size for different $C_d$ and $f_A$. For each combination of $C_d \in [0, 3]$ and $f_A \in [0.05, 0.95]$ by step size 0.05, we computed $p_A$ and $p_B$ and ran the SIR dynamics as described in the main text. We set $E_d = 0.25$. We then calculated the final epidemic size as $1 - \frac{S_t}{S_0}$ where $S_0$ and $S_t$ are the number of susceptible individuals at the beginning and end of the epidemic respectively.

We found that the final epidemic size depends on both $C_d$ and $f_A$ (Fig S9). For a specific $f_A$, epidemic size decreases as $C_d$ increases. This was expected as higher $C_d$, equivalently, more heterogeneity in susceptibility, is known to result in smaller epidemics (Ball, 1985). However, for a specific $C_d$, there can be different epidemic sizes depending on $f_A$, and if both $f_A$ and $C_d$ are changed, it is possible to get a larger epidemic with a higher $C_d$. The smallest epidemic is possible when $f_A = E_d$. This is because, when $f_A = E_d$, $p_A \to 1$ and $p_B \to 0$ as $C_d$ increases. So, with a large $C_d$, the largest possible final epidemic size here occurs when all the $A$ individuals are infected. This gives an epidemic size of $E_d$. When $f_A$ shifts away from $E_d$, $p_A$ remains large but $p_B > 0$, so the epidemic can continue even if all $A$ individuals are infected. This can then result in an epidemic size greater than $E_d$. Overall, in addition to $C_d$ and $E_d$, the fraction of the population that is the more susceptible type of individual $f_A$ is critical for determining the trajectory of the epidemic.

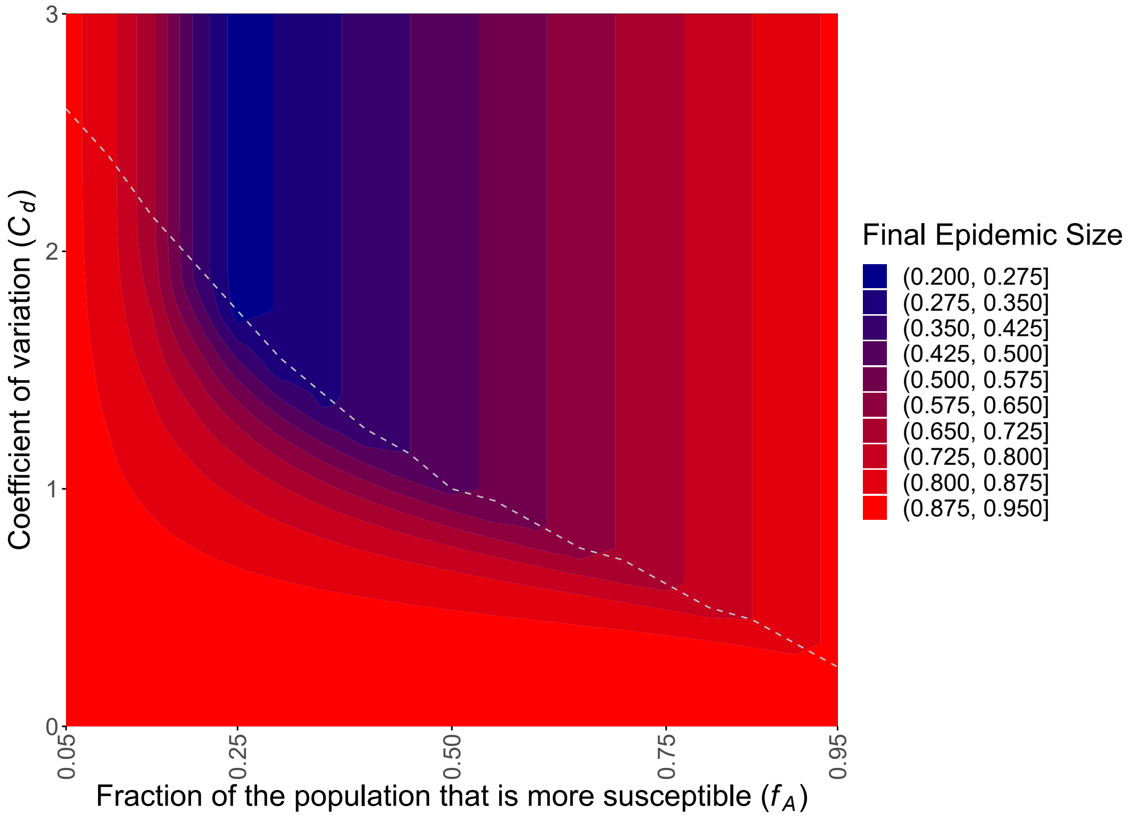


Figure S9: Final epidemic size in the discrete case depends on both the coefficient of variation of risk $C_d$ and fraction of the population that is more susceptible $f_A$. The plot shows the fraction of individuals infected over the course of an epidemic with varying $C_d$ and $f_A$. The area above the gray dashed line represents parameter space that gives computationally indistinguishable probabilities of infection $p_A$ and $p_B$, and therefore final epidemic size, to the parameter combination with the same $f_A$ and highest $C_d$ below the line. This occurs because risks of infection can be changed to increase $C_d$ without bound, whereas probabilities are bounded. Note that for a specific $f_A$, epidemic size decreases as $C_d$ increases, but for a specific $C_d$, epidemic size differs depending on $f_A$. The smallest epidemic is possible when $f_A = E_d$. $E_d = 0.25$.

### S7. Effect of having informative priors for $p_A$, $p_B$, or $f_A$ on predicted disease dynamics

In the discrete case, we are unable to generate as precise of disease dynamics predictions as in the continuous case. This is because in the discrete case, we must estimate the three parameters $p_A$, $p_B$, and $f_A$ to define the risk distribution, whereas in the continuous case, only two parameters $k$ and $\theta$ must be estimated. In addition, the parameters are all highly correlated, leading to low identifiability. If, however, anything is previously known about the discrete parameters, then we may be able to more precisely estimate them by setting informative priors. For example, if the factor creating heterogeneity in susceptibility is whether someone wears a mask or not, we might have an idea of $f_A$ (i.e., the fraction of people that do not wear masks). To test this, we generated posterior distributions using Metropolis-Hastings MCMC and ABC as described in the main text but with informative priors instead of flat priors. For the priors, we used truncated normal distributions restricted to $[0, 1]$ with mean given by the true parameter value for $p_A$, $p_B$, or $f_A$ and standard deviation 0.05. We then randomly sampled $1,000$ parameter sets from the posterior distribution to run SIR model simulations and determine 95% CIs. Unsurprisingly, we found that we could generate narrower 95% CIs with informative priors than with flat priors (Fig S10). This indicates that prior knowledge of $p_A$, $p_B$, or $f_A$ could improve the precision of disease dynamics predictions in the discrete case.

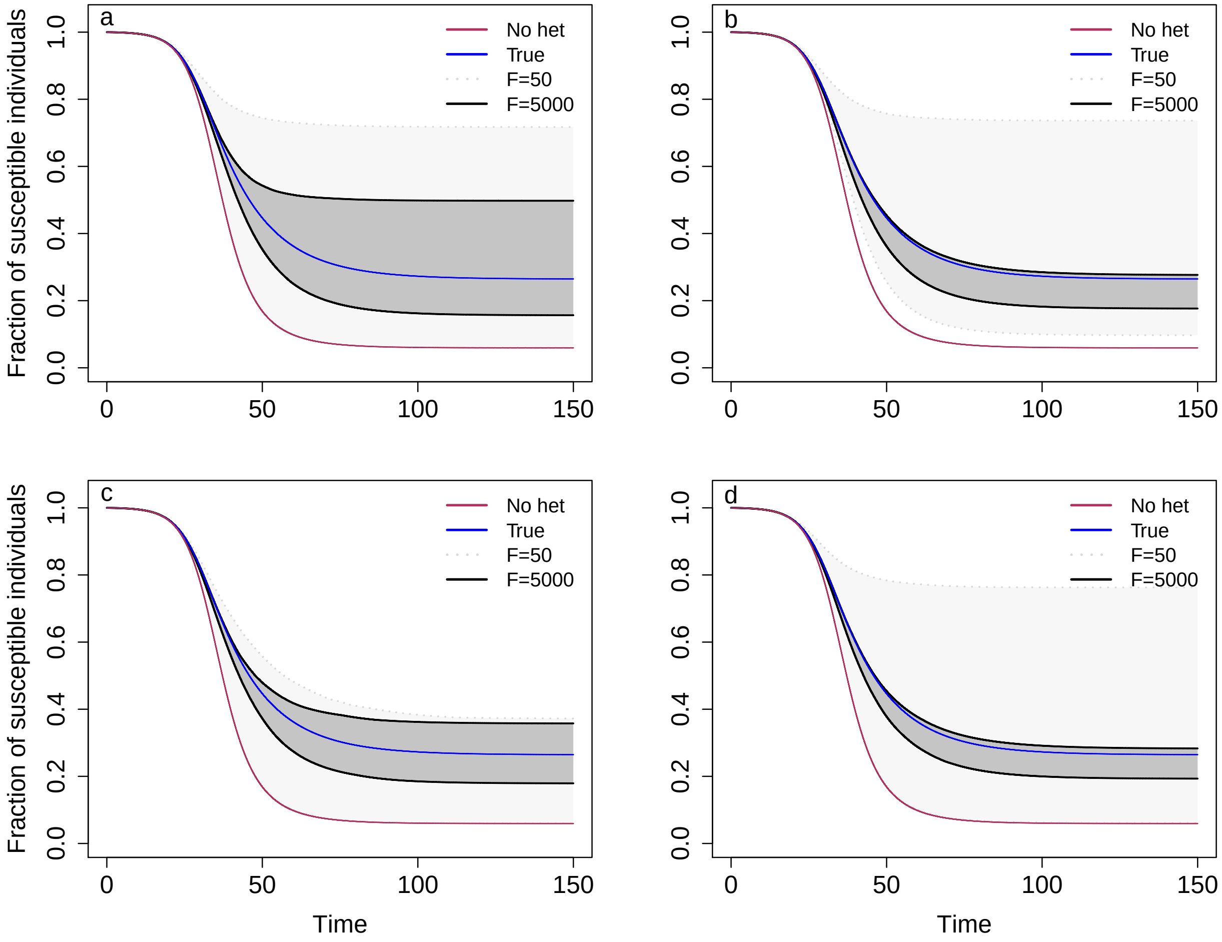


Figure S10: Using informative priors for $p_A$, $p_B$, or $f_A$ improves the precision of our predicted disease dynamics. The plots show the predicted SIR dynamics in the discrete case with a) uninformative priors for $p_A$, $p_B$, and $f_A$, b) an informative prior for $p_A$, c) an informative prior for $p_B$, and d) an informative prior for $f_A$. Specifically, the fraction of susceptible individuals $\frac{S}{S_0}$ is shown over the course of an epidemic. Shaded regions represent 95% CIs determined from 1,000 posterior samples for $F = 50$ (light gray), 200 (medium gray), 1000 (dark gray), and 5000 (black). The blue line shows the true dynamics for the parameters used to generate the contact tracing data, and the red line shows the corresponding dynamics if there is homogeneity in susceptibility. The informative priors used for $p_A$, $p_B$, and $f_A$ were truncated normal distributions restricted to [0,1] with mean given by the true parameter value and standard deviation 0.05. $C_d = 1.3$, $E_d = 0.25$, $f_A = 0.2$, and $N = 5$.

### S8. False negatives

While in the main text we have assumed access to perfect data, we can imagine that real contact tracing data may be imperfect. One way we have considered that data may be imperfect is that individuals may be mislabeled as uninfected when they are infected (false negatives). As we said in the main text, false negatives can cause our method to overestimate the level of heterogeneity in susceptibility because our estimate of the infection probability for focal individuals $p_f$ may be biased lower. This is because, assuming infection confers at least partial immunity, focal individuals that were previously infected (i.e. false negatives) will be less likely to be infected than focal individuals that were true negatives. To counteract this issue, we developed a version of the method that corrects for false negatives by adjusting the likelihood calculations for both detecting and estimating heterogeneity in susceptibility. For detection, we first consider that $x_I$ of the $F$ focal individuals were false negatives and therefore incorrectly classified as focal with probability $p_I$, so we should only include $F - x_I$ focal individuals in the likelihood. Then, if we observe $D_N$ naive individuals and $D_F$ focal individuals infected in the data, there may be an additional $x_N$ naive individuals and $x_F$ focal individuals infected that we observed as uninfected where $x_N \in [0, F(N-1) - D_N]$ and $x_F \in [0, F - x_I - D_F]$. We changed the likelihood equations to the following in order to account for false negatives.

$$
\begin{aligned}
L_{\text{hom}} =& \ln\left[\sum_{x_N=0}^{F(N-1)-D_N}\left[P(D_N + x_N|F(N-1), \bar{p}) * P(x_N|D_N + x_N, \beta)\right]\right] \\
&+ \ln\left[\sum_{x_I=0}^{F}\sum_{x_F=0}^{F-x_I-D_F}\left[P(D_F + x_F|F - x_I, \bar{p}) * P(x_I|F, p_{I,\text{hom}}) * P(x_F|D_F + x_F, \beta)\right]\right] \qquad (1)
\end{aligned}
$$

$$
\begin{aligned}
L_{\text{het}} =& \ln\left[\sum_{x_N=0}^{F(N-1)-D_N}\left[P(D_N + x_N|F(N-1), p_n) * P(x_N|D_N + x_N, \beta)\right]\right] \\
&+ \ln\left[\sum_{x_I=0}^{F}\sum_{x_F=0}^{F-x_I-D_F}\left[P(D_F + x_F|F - x_I, p_f) * P(x_I|F, p_{I,\text{het}}) * P(x_F|D_F + x_F, \beta)\right]\right] \qquad (2)
\end{aligned}
$$

$L_{\text{hom}}$ is the log-likelihood of the data under the null hypothesis that there is homogeneity in susceptibility, and $L_{\text{het}}$ is the log-likelihood under the alternative hypothesis that there is heterogeneity in susceptibility. $P(x|n,p)$ is the probability of observing $x$ individuals infected out of $n$ individuals exposed with probability $p$ of being infected and is distributed according to a binomial distribution. $D_N + x_N$ and $D_F + x_F$ are the numbers of naive and focal individuals actually infected respectively where $D_N$ is the number of naive individuals observed infected in the data, $D_F$ is the number of focal individuals observed infected in the data, and $x_N$ and $x_F$ are the numbers of naive and focal individuals infected but observed as uninfected (number of false negatives). $D_N + x_N \in [0, F(N-1)]$, $x_N \in [0, F(N-1) - D_N]$, $D_F + x_F \in [0, F - x_I]$, and $x_F \in [0, F - x_I - D_F]$. $x_I$ is the number of individuals that were previously infected and incorrectly classified as focal because they were perceived to be uninfected. $x_I \in [0, F]$. $\beta$ is the false negative rate where $\beta \in [0, 1]$. $p_{I,\text{hom}}$ and $p_{I,\text{het}}$ are the probabilities that a focal individual was previously infected and incorrectly classified for the homogeneous and heterogeneous likelihoods respectively. $p_{I,\text{hom}} = \frac{\bar{p}\beta}{1-\bar{p}+\bar{p}\beta}$ and $p_{I,\text{het}} = \frac{p_n\beta}{1-p_n+p_n\beta}$. $p_n$, $p_f$ and $\bar{p}$ are the true probabilities of infection and are calculated as $p_n = \frac{p_{n,\text{obs}}}{1-\beta}$, $p_f = \frac{p_{f,\text{obs}}F}{(F-x_I)(1-\beta)}$, and $\bar{p} = \frac{\bar{p}_{\text{obs}}FN}{(FN-x_I)(1-\beta)}$. $p_{n,\text{obs}}$, $p_{f,\text{obs}}$ and $\bar{p}_{\text{obs}}$ are the probabilities of infection observed from the data including false negatives and are estimated as $p_{n,\text{obs}} = \frac{D_N}{F(N-1)}$, $p_{f,\text{obs}} = \frac{D_F}{F}$, and $\bar{p}_{\text{obs}} = \frac{D_F+D_N}{FN}$.

For estimation, we also altered the likelihood calculations in our MCMC algorithm. In the discrete case, we implemented the same ABC framework as in the main text except we included false negatives at rate $\beta$ in our simulations. In the continuous case, we switched from an analytical likelihood to a simulated likelihood with ABC. We also increased our MCMC chain to length 1,200,000 with a burn-in of 500,000 and thinning interval 100. We followed the same procedure to calculate likelihood in the continuous case as in the discrete case.

For detecting heterogeneity in susceptibility, adjusting the likelihood calculation greatly lessened the impact of false negatives, but the method will still fail for values of $E$ close to 1 (Figs S11, S12). This is because our likelihoods are approximations as we do not account for stochasticity in the observed number of infected individuals and the number of false negatives in our calculations. This means that the true probabilities of infection will be computed as greater than 1 if the observed probability of infection and false negative rate $\beta$ are large enough. For example, if $p_{n,\text{obs}} > 1 - \beta$, then $p_n > 1$. Since probabilities cannot exceed 1, we set an upper bound of 1 for the true probabilities. However, if there are no false negatives in the data, we could incorrectly compute a true probability of infection as 1 when it is actually less than 1. For values of $E_{\text{inf}}$ close to 1, this can cause us to incorrectly detect or not detect heterogeneity in susceptibility. We do not think this will be a major issue though as $E$ is typically less than 0.5 (Ajelli et al., 2015; Koh et al., 2020; Lessler et al., 2009; Taylor et al., 2007; De Serres et al., 2000; Rieder, 2003). For estimating parameters and predicting disease dynamics, adjusting the likelihoods to correct for false negatives fixed the issue. This is shown in Figure S13 where the 95% CI for the method adjusted to account for false negatives (black) captures the true disease dynamics, whereas the original, unadjusted method (gray) overestimates the level of heterogeneity, leading to the prediction of a smaller epidemic.

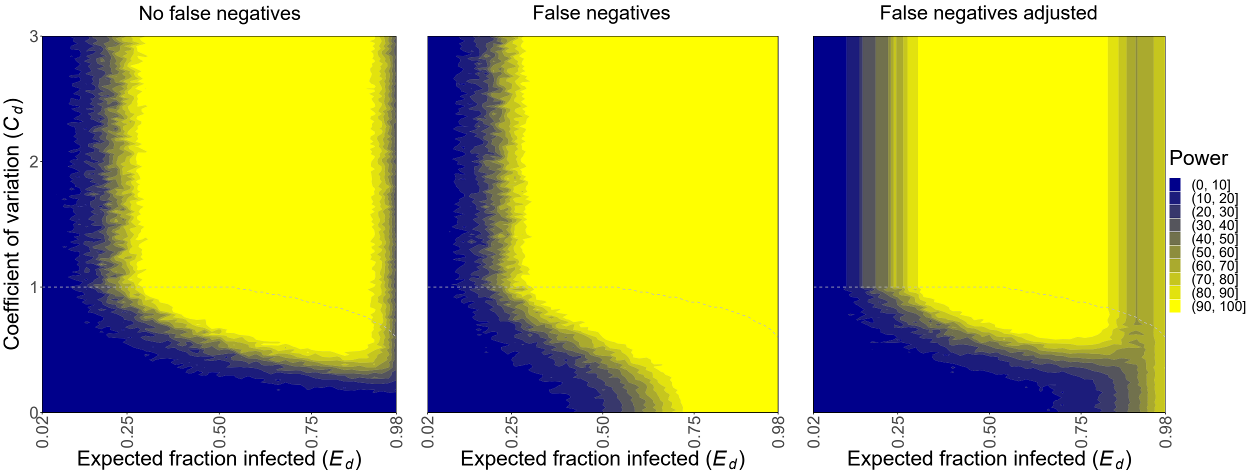


Figure S11: The presence of false negatives in the discrete case causes our method to detect heterogeneity in susceptibility when we should not be able to, but adjusting the method corrects this issue except when $E_d$ is close to 1. The plots show the power to detect heterogeneity in susceptibility in the discrete case with no false negatives, false negatives with the original method, and false negatives with the adjusted method. The areas above the gray dashed lines represent parameter space that gives computationally indistinguishable probabilities of infection $p_A$ and $p_B$, and therefore power, to the parameter combination with the same $E_d$ and highest $C_d$ below the line. This occurs because risks of infection can be changed to increase $C_d$ without bound, whereas probabilities are bounded. $F = 200$, $N = 5$, $f_A = 0.5$, and $\beta = 0.1$.

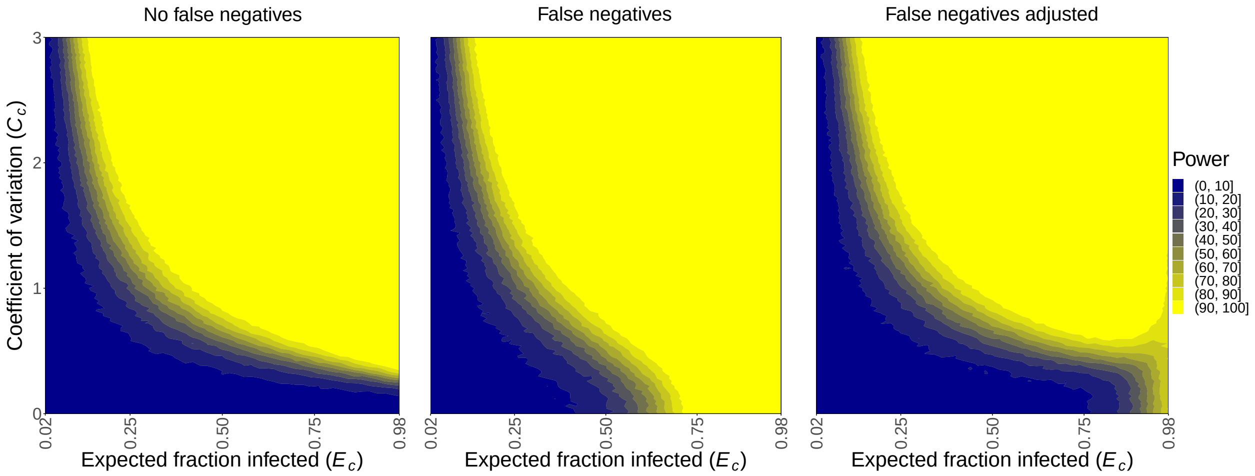


Figure S12: The presence of false negatives in the continuous case causes our method to detect heterogeneity in susceptibility when we should not be able to, but adjusting the method corrects this issue except when $E_c$ is close to 1. The plots show our power to detect heterogeneity in susceptibility in the continuous case with no false negatives, false negatives with the original method, and false negatives with the adjusted method. $F = 200$ and $\beta = 0.1$.

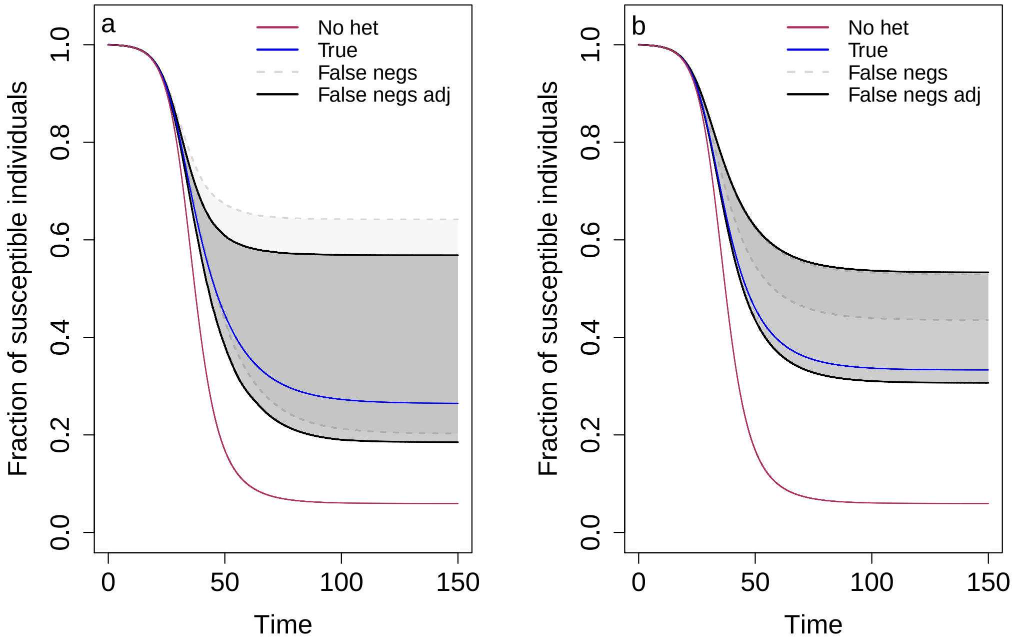


Figure S13: The presence of false negatives causes our method to overestimate the level of heterogeneity in susceptibility, but adjusting the method corrects this issue. The plots show the predicted SIR dynamics in a) the discrete case and b) the continuous case with false negatives when we do or do not adjust the method to account for false negatives. Specifically, the fraction of susceptible individuals $\frac{S}{S_0}$ is shown over the course of an epidemic. Shaded regions represent 95% CIs determined from 1,000 posterior samples for the method adjusted to account for false negatives (black) and not adjusted to account for false negatives (gray). The blue line shows the true dynamics for the parameters used to generate the contact tracing data, and the red line shows the corresponding dynamics if there is homogeneity in susceptibility. $C_d = 1.3$, $E_d = 0.25$, $f_A = 0.2$, $\beta_d = 0.2$, $C_c = 1$, $E_c = 0.75$, $\beta_c = 0.1$, $F = 1000$, and $N = 5$.

### S9. Heterogeneity in transmission

As mentioned in the main text, heterogeneity in transmission, differences between individuals in their likelihood of infecting others, given contact may create problems for our method. This is because heterogeneity in transmission given contact may swamp out perceived differences in individuals' susceptibilities as force of infection in addition to innate susceptibility affect whether individuals become infected, so less susceptible individuals are not necessarily less likely to be infected. To test this, we attempted to detect and estimate heterogeneity in susceptibility given contact as well as predict SIR dynamics in the presence of heterogeneity in transmission given contact. We used the same methods as in the main text but simulated the data with heterogeneity in transmission in addition to heterogeneity in susceptibility. In order to add heterogeneity in transmission into our simulations, we generated a force of infection $\lambda_i$, $i = 1, ..., 2F$ to represent the infectiousness of the infected individual in each contact network where $\lambda_i$ has distribution Gamma$(m, \phi)$ with mean $m\phi = 1$. The shape parameter $m$ is a measure of the level of heterogeneity in transmission present where small $m$ means that there is a lot of heterogeneity, and as $m \to \infty$, the level of heterogeneity decreases to zero. Here, we set $m = 0.5$ and $\phi = 2$ as this falls within the range of biologically reasonable shape parameters (Lloyd-Smith et al., 2005). We chose the gamma distribution because it is flexible and is used as part of the Poisson-gamma mixture definition of the negative binomial distribution to describe the number of individuals infected by a particular infected individuals (Lloyd-Smith et al., 2005). We then multiplied the force of infection by the risk of being infected given contact to compute the probability of infection for each individual as $p_{i,j} = 1 - e^{-\lambda_i r_j}$ for $j = A, B$ in the discrete case and $j = 1, ..., FN$ in the continuous case. We also calculated an average force of infection $\Lambda$ across all networks to ensure that we kept the same average probability of infection at the beginning of the epidemic regardless of heterogeneity. This was also multiplied by the force of infection for each network and risk of being infected to give the probability of infection for each individual as $p_{i,j} = 1 - e^{-\lambda_i \Lambda r_j}$ for $j = A, B$ in the discrete case and $j = 1, ..., FN$ in the continuous case. We computed $\Lambda$ by numerically solving for the $\Lambda$ that made it so that the expected probability of infection without heterogeneity in transmission, $\mathrm{E}[1 - e^{-r_j}]$, equaled the expected probability of infection with heterogeneity in transmission, $\mathrm{E}[1 - e^{-\lambda_i \Lambda r_j}]$.

We found that our power to detect heterogeneity in susceptibility and our ability to predict SIR dynamics in the presence of heterogeneity in transmission were impacted considerably (Figs S14, S15, S16). With heterogeneity in transmission given contact, our power was greatly reduced, and we made less precise predictions of the disease dynamics. We also predicted a larger final epidemic size in the continuous case as we underestimated the level of heterogeneity in susceptibility. This is because our method does not consider that the force of infection in each contact network affects whether individuals are infected in addition to their innate susceptibilities. Our method, in its current form, is therefore not suitable for these cases. However, even though our power was reduced, we note that the presence of heterogeneity in transmission did not cause us to detect heterogeneity in susceptibility when it did not exist (i.e. $C = 0$). Additionally, the impact of heterogeneity in transmission would be smaller with a larger $m$ (i.e. less heterogeneity in transmission) and non-existent if this heterogeneity was due to contact rate rather than innate differences. Heterogeneity in contact rate would simply mean that each network would have its own number of contacts $N$. Importantly, we note that this variation in contact rate is typically assumed to be the cause of heterogeneity in transmission (Lloyd-Smith et al., 2005) in which case our method would be suitable.

One solution to manage the effect of heterogeneity in transmission given contact on our method is to test for its presence. This allows us to determine whether the assumption of our model that there is no heterogeneity in transmission is violated. If there is not heterogeneity in transmission given contact, the number of naive individuals infected in each contact network will be binomially distributed regardless of the presence of heterogeneity in susceptibility given contact. This is because individuals are as likely to be infected or not if they are in contact with different hosts. And, when there is heterogeneity in susceptibility, individuals with higher and lower susceptibilities are binomially distributed across networks. If there is heterogeneity in transmission, we would expect to see a disproportionate number of contact networks where most or few individuals are infected. In the extreme case, either everyone or no one will be infected in each network. This is because the force of infection in each network will strongly affect whether individuals in that network are infected.

To test for heterogeneity in transmission given contact, we can perform a goodness of fit test that compares the distribution of infection in naive individuals among contact networks to see if it follows a single binomial

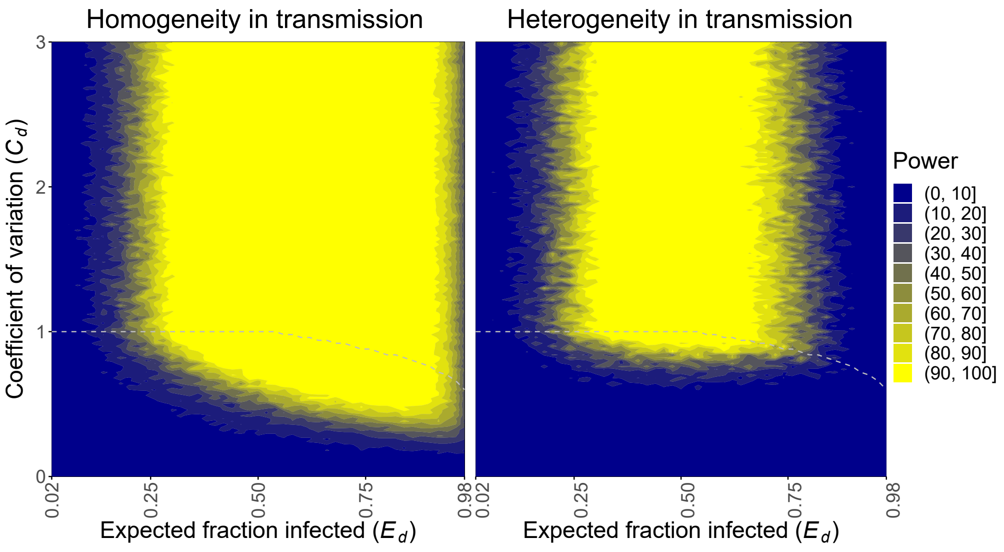


Figure S14: The presence of heterogeneity in transmission given contact reduces our power to detect heterogeneity in susceptibility in the discrete case. The plots show the power to detect heterogeneity in susceptibility in the discrete case with homogeneity in transmission and with heterogeneity in transmission. The areas above the gray dashed lines represent parameter space that gives computationally indistinguishable probabilities of infection $p_A$ and $p_B$, and therefore power, to the parameter combination with the same $E_d$ and highest $C_d$ below the line. This occurs because risks of infection can be changed to increase $C_d$ without bound, whereas probabilities are bounded. $F = 200$, $N = 5$, $f_A = 0.5$, $m = 0.5$, and $\phi = 2$.

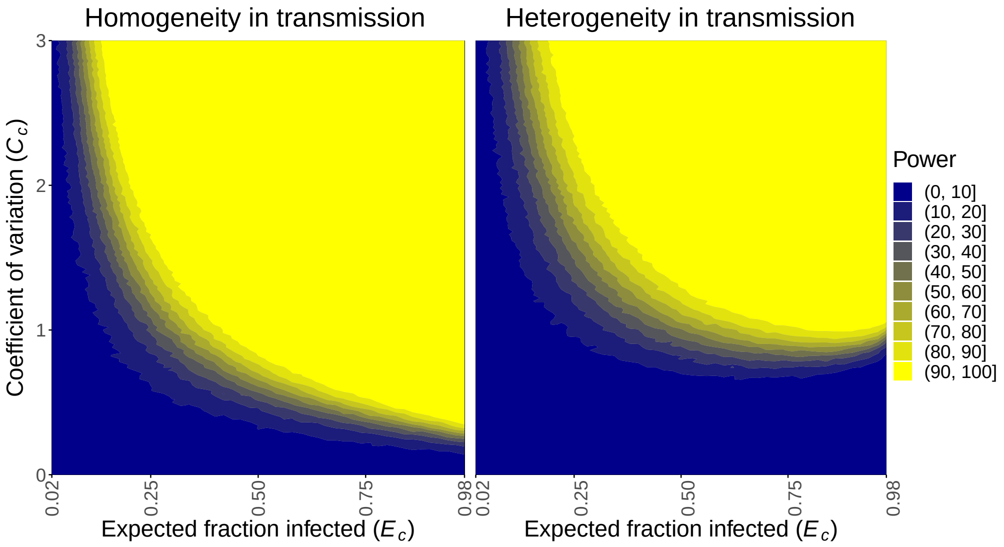


Figure S15: The presence of heterogeneity in transmission given contact reduces our power to detect heterogeneity in susceptibility in the continuous case. The plots show the power to detect heterogeneity in susceptibility in the continuous case with homogeneity in transmission and with heterogeneity in transmission. $F = 200$, $N = 5$, $m = 0.5$, and $\phi = 2$.

distribution. We use only naive individuals so that there is no effect on average susceptibility from focal individuals in case there is heterogeneity in susceptibility. To do so, we essentially simulate the contact networks many times with a binomial distribution and compare the likelihoods of those data to the real data to check if the real data are consistent with homogeneity in transmission. We first calculate an overall probability of infection $p_n$ across all networks as the number of naive individuals infected divided by the number of naive individuals exposed. Next, we simulate the number of naive individuals infected in each network $10,000$ times according to a binomial distribution with probability $p_n$ of infection. We can then

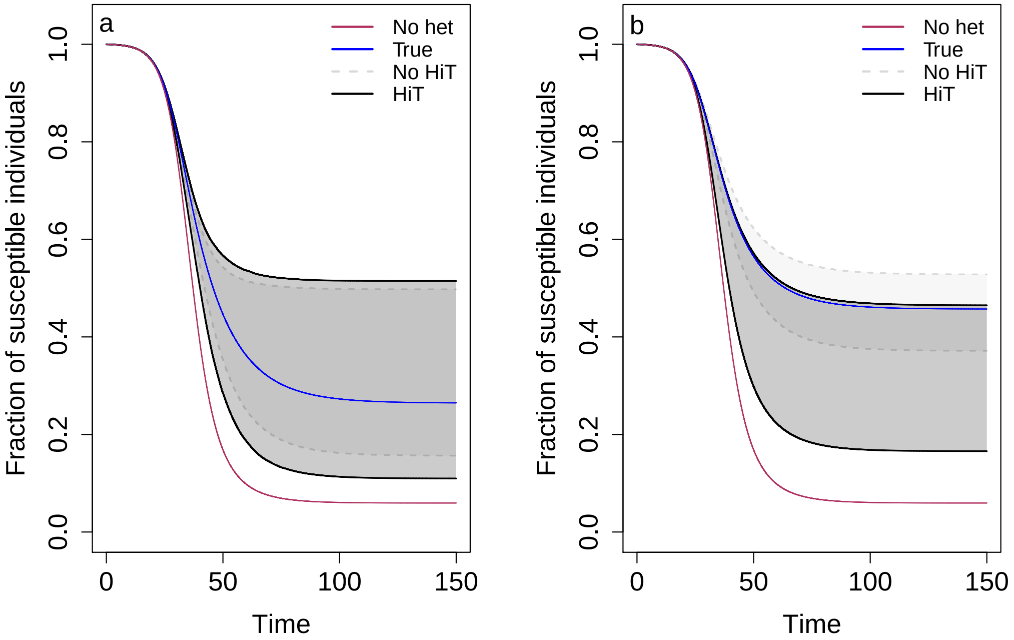


Figure S16: The presence of heterogeneity in transmission given contact decreases the precision of our prediction of the disease dynamics and can cause our method to underestimate the level of heterogeneity in susceptibility. The plots show the effect of heterogeneity in transmission (HiT) on predicted SIR dynamics in a) the discrete case and b) the continuous case. Specifically, the fraction of susceptible individuals $\frac{S}{S_0}$ is shown over the course of an epidemic. Shaded regions represent 95% CIs determined from 1,000 posterior samples for homogeneity in transmission (gray) and heterogeneity in transmission (black). The blue line shows the true dynamics for the parameters used to generate the contact tracing data, and the red line shows the corresponding dynamics if there is homogeneity in susceptibility. $C_d = C_c = 1.3$, $E_d = E_c = 0.25$, $f_A = 0.2$, $F = 5000$, $N = 5$, $m = 0.5$, and $\phi = 2$.

calculate the log-likelihood of the real data as well as the log-likelihood of each set of simulated data as $\sum_{i=1}^{F} \ln[P(x_i|N_i, p_n)]$. $x_i$ is the number of naive individuals infected in network $i$ and $N_i$ is the number of naive individuals exposed in network $i$. If the log-likelihood of the real data is within the 95% CI of the simulated log-likelihoods, we conclude that there is not significant heterogeneity in transmission given contact. If not, we conclude that there is heterogeneity in transmission given contact, and, therefore, our method to detect heterogeneity in susceptibility would be underpowered, and estimates of heterogeneity in susceptibility would be unreliable (Fig S17).

We tested the ability of this method to detect heterogeneity in transmission for parameters of the distribution dictating the level of heterogeneity in transmission $m \in [0.1, 6]$ with step size 0.1 and $\phi = \frac{1}{m}$. We examined $m \in [0.1, 6]$ because this captures a range of published estimates for this parameter: 0.16 to 5.1 (Lloyd-Smith et al., 2005). We also set $C_d = C_c = 1.3$, $E_d = E_c = 0.25$, $f_A = 0.2$, $N = 5$, and $F = 200$ or 1000. This was done for $1,000$ simulations to compute the statistical power of the method. Our power here is the percent of simulations in which the log-likelihood of the real data lies outside the 95% CI of the simulated log-likelihoods (i.e. we detect heterogeneity in transmission given contact).

Figure S18 shows that the sample size and level of heterogeneity affect our power to detect heterogeneity in transmission. As the number of focal individuals $F$ increases from 200 to 1000 and as the number of contacts per network $N$ increases from 5 to 10, there is greater power to detect all levels of heterogeneity in transmission ($m$). This was to be expected because a higher sample size improves the precision of our likelihoods. The level of heterogeneity in transmission present is described by the shape parameter $m$ of the distribution of forces of infection. As $m$ decreases, there is more power to detect heterogeneity in transmission as there is more heterogeneity in the population.

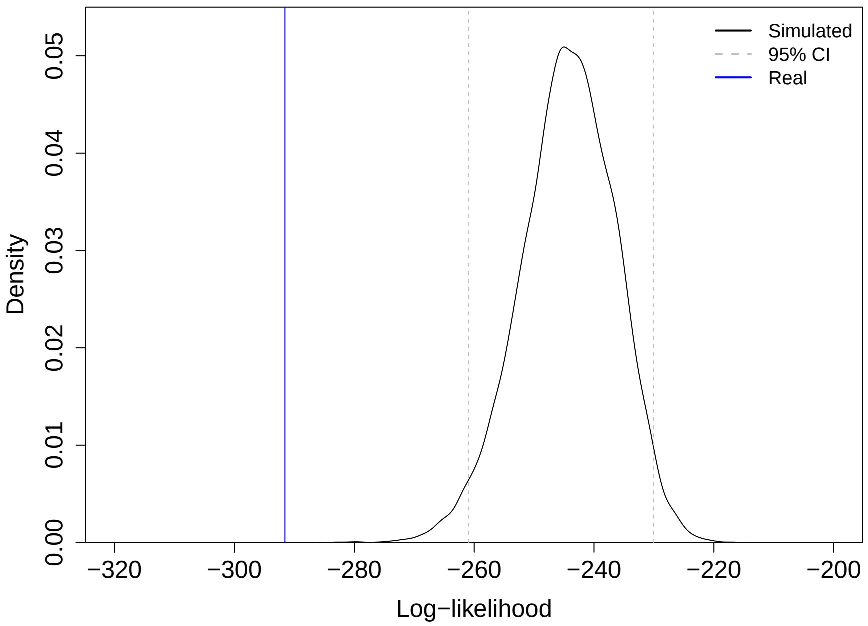


Figure S17: Example of the method to detect heterogeneity in transmission given contact. The black line shows the distribution of simulated log-likelihoods, the gray dashed lines show the 95% CI of the simulated log-likelihoods, and the blue line shows the log-likelihood of the real data. Here, we would conclude that there is heterogeneity in transmission as the log-likelihood of the real data falls outside the 95% CI of the simulated log-likelihoods. $C_d = 1.3$, $E_d = 0.25$, $f_A = 0.2$, $F = 200$, $N = 5$, $m = 0.5$, and $\phi = 2$.

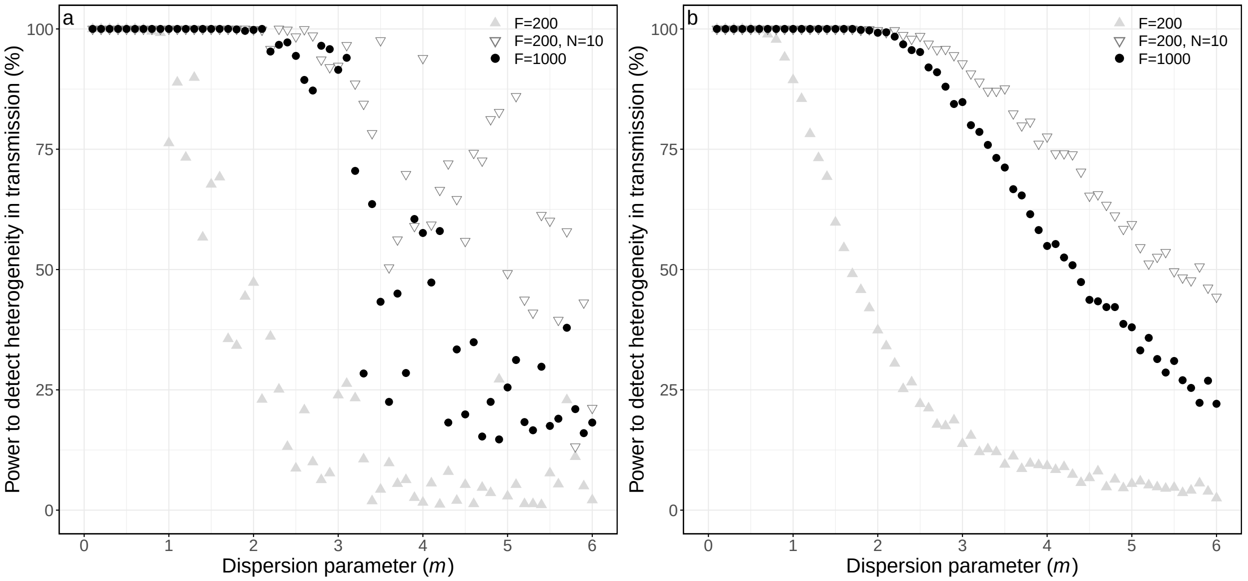


Figure S18: An increased sample size (larger $F$ and $N$) and increased level of heterogeneity in transmission given contact (smaller $m$) enhance our power to detect heterogeneity in transmission given contact. The plots show the power to detect heterogeneity in transmission given contact in the a) discrete case and b) continuous case in the presence of heterogeneity in susceptibility across varying levels of heterogeneity in transmission. Specifically, they show the percent of simulations in which the distribution of infection in naive individuals among contact networks is not binomial, so we conclude that there is heterogeneity in transmission given contact for $F = 200$ and $N = 5$ (light gray triangles), $F = 200$ and $N = 10$ (dark gray triangles), or $F = 1000$ and $N = 5$ (black circles). Heterogeneity in transmission is gamma distributed with shape parameter $m$ and scale parameter $\phi = \frac{1}{m}$. $C_d = C_c = 1.3$, $E_d = E_c = 0.25$, and $f_A = 0.2$.
